# TNFR1-mediated senescence and lack of TNFR2-signaling limit human intervertebral disc cell repair in back pain conditions

**DOI:** 10.1101/2024.02.22.581620

**Authors:** Jennifer Gansau, Elena Grossi, Levon Rodriguez, Minghui Wang, Damien M. Laudier, Saad Chaudhary, Andrew C. Hecht, Wenyu Fu, Robert Sebra, Chuanju Liu, James C. Iatridis

**Affiliations:** Department of Orthopaedics, Icahn School of Medicine at Mount Sinai; New York, NY 10029, USA; Department of Oncological Sciences, Icahn School of Medicine at Mount Sinai; New York, NY 10029, USA; Department of Dermatology, Icahn School of Medicine at Mount Sinai; New York, NY 10029, USA; Department of Genetics & Genomic Sciences, Icahn School of Medicine at Mount Sinai; New York, NY 10029, USA; Department of Orthopaedics & Rehabilitation, Yale University School of Medicine; New Haven, CT 06510, USA

## Abstract

Poor intervertebral disc (IVD) healing causes IVD degeneration (IVDD) and progression to herniation and back pain. This study identified distinct roles of TNFα-receptors (TNFRs) in contributing to poor healing in painful IVDD. We first isolated IVDD tissue of back pain subjects and determined the complex pro-inflammatory mixture contained many chemokines for recruiting inflammatory cells. Single-cell RNA-sequencing of human IVDD tissues revealed these pro-inflammatory cytokines were dominantly expressed by a small macrophage-population. Human annulus fibrosus (hAF) cells treated with IVDD-conditioned media (CM) underwent senescence with greatly reduced metabolic rates and limited inflammatory responses. TNFR1 inhibition partially restored hAF cell metabolism sufficiently to enable a robust chemokine and cytokine response to CM. We showed that the pro-reparative TNFR2 was very limited on hIVD cell membranes so that TNFR2 inhibition with blocking antibodies or activation using Atsttrin had no effect on hAF cells with CM challenge. However, TNFR2 was expressed in high levels on macrophages identified in scRNA-seq analyses, suggesting their role in repair responses. Results therefore point to therapeutic strategies for painful IVDD involving immunomodulation of TNFR1 signaling in IVD cells to enhance metabolism and enable a more robust inflammatory response including recruitment or delivery of TNFR2 expressing immune cells to enhance IVD repair.

**SUMMARY STATEMENT:** TNFR1 signaling drives cells towards senesce and muted inflammatory response in painful intervertebral disc degeneration, while limited TNFR2 signaling may limit disc cell repair responses.

## INTRODUCTION

Back and neck pain are leading causes of global disability impacting >100 million US adults, leading to missed workdays, massive health care expenditures and contribute to opioid usage, and represents a tremendous global socioeconomical burden (1–4). Intervertebral disc (IVD) herniation involving annulus fibrosus (AF) defects is a direct cause of pain and disability that can predispose to IVD degeneration (IVDD) and result in painful disability (1, 5, 6). Chronic inflammation is a characterizing feature of IVDD and herniation with pro-inflammatory cytokines contributing to progressive IVDD and pain (7–10). Pro-inflammatory cytokines play known roles in IVD matrix catabolism, neurovascular ingrowth, nociception and cellular senescence (8–20). Tumor necrosis factor-alpha (TNFα), interleukin-1 Beta (IL-1β) and interleukin-6 (IL-6) are major cytokines identified in the IVD known to propagate matrix catabolism and nociception, and considered targets for therapeutic strategies (8, 9, 11–14, 20, 21).

Human clinical trials and animal studies show promise for TNFα blocking in IVDD, but mixed results prompt a need for further research. In human clinical trials for IVD herniation with radiculopathy, local and systemic administration of TNFα blocking agents, including infliximab, etanercept and adalimumab, showed some promise for pain reduction but remained inconclusive since trials exhibited broad heterogeneity in effects and several showed negative findings (22–28). While a randomized, double-blinded, placebo-controlled trial of 43 patients with disc herniation found some decrease of average back pain using etanercept compared to placebo after 28 weeks, a randomized, controlled trial that enrolled 40 discectomy patients with disc herniation concordant with radicular pain, found infliximab had similar effects as placebo at 3-month follow-up (23, 29). Meanwhile, preclinical animal studies show efficacy with early TNFα blocking with reduced pain and improved IVD degeneration score (13, 30). These mixed clinical and animal results highlight importance of TNFα, but also a need to better understand TNFα signaling and crosstalk between IVD and immune cells (7). Therefore, here, we focus on specific roles of TNFα-receptors.

TNFα signaling occurs via 2 receptors: TNFα-receptor 1 (TNFR1) is implicated in the pro-inflammatory response that promote tissue catabolism, and TNFα-receptor 2 (TNFR2) is associated with cell survival and tissue repair/regeneration (31–34). Current treatment strategies involve antibody-based inhibition with a high binding affinity for TNFα and therefore actively prevents TNFα from binding to both TNFR1 and TNFR2 (23, 30, 32, 35, 36). Selective TNFR inhibition shows promise for IVDD since TNFR1^-/-^ mice protected from IVDD while TNFR2^-/-^ mice accelerated IVDD (20). The engineered protein Atsttrin is a TNFR2 activator derived from progranulin, and this novel therapeutic was shown in osteoarthritis and IVD organ culture to shift TNFα signaling from TNFR1 to a greater TNFR2 response (37–41). However, these studies need validation in a human model system with a translational strategy.

A common strategy for simulating IVDD conditions is supplementation of cell culture media with pro-inflammatory cytokines such as TNFα, IL-1β and IL-6 (10, 37, 42–44). However, painful human IVDD with herniation is known to involve multiple cytokines and chemokines that are collectively responsible for different cellular functions thought to inhibit IVD repair and promote painful responses (8). We therefore applied a human relevant cell culture model system that recapitulated the complex cytokine environment of human IVDD.

This study tested the hypotheses that human IVDD involves a complex cytokine mixture that drives IVD cells towards senescence and can be shifted towards a reparative responsive by inhibiting TNFR1 and promoting TNFR2 activity. This study used human tissue and cells from painful IVDD subjects and autopsies, and applied cell culture, bulk RNA-sequencing, single-cell RNA-sequencing (scRNA-seq), gene and protein assays. We determined that the many cytokines and chemokines present in IVDD tissue from surgical subjects were dominantly expressed by a small population of macrophages, and expressed at far lower levels by IVD cells. Next, we showed that IVDD conditioned media (CM) treatment on human AF (hAF) cells *in vitro* had the largest effects inhibiting metabolic rates and promoting senescence. Blocking TNFR1 signaling during CM exposure sufficiently restored hAF cell metabolism to enable a robust pro-inflammatory response. However, blocking TNFR2 and enhancing TNFR2 signaling with Atsttrin during CM exposure had no effects on hAF cells. Finally, we showed TNFR1 is highly expressed by IVD cells and increased with IVDD grade underscoring its importance in IVDD, and we showed that TNFR2 was expressed at minimal amounts by IVD cells suggesting limited TNFR2 signaling at below therapeutic levels in IVDD conditions.

## RESULTS

### CM contains many cytokines & chemokines dominantly expressed by macrophages

To characterize the IVDD pro-inflammatory microenvironment, human surgical IVD specimens containing both AF and NP portions were collected (Table S1) and cultured in Basal media for 72 hours under constant agitation, leading to cytokine release and generating IVDD derived CM (Fig. 1A). CM cytokines and chemokines were analyzed with a human cytokine/chemokine panel 48-plex discovery Assay® Array, with pro-inflammatory cytokines (e.g., IL-1β, IL-6, TNFα), anti-inflammatory cytokines (e.g., IL-1RA, IL-4, IL-10, IL-13), and chemokines (e.g., MIP-1α/CCL3, MIP1β/CCL4, MCP-1/CCL2 and MCP-3/CCL7) detected (Fig. 1B, Table S2). To determine the cells producing these factors, we performed scRNA-seq on cells isolated from 5 herniated IVD specimens containing AF and NP regions from 3 subjects undergoing Anterior cervical discectomy and fusion (ADF) procedure (1 male, 2 female) (Fig. 1A, Fig. S1A, Table S1). The 13,259 cells were analyzed to identify 14 clusters that were annotated using UniCell Deconvolve (Fig. S1), a pre-trained, unbiased, deep learning model system (45).

**Fig. 1.**
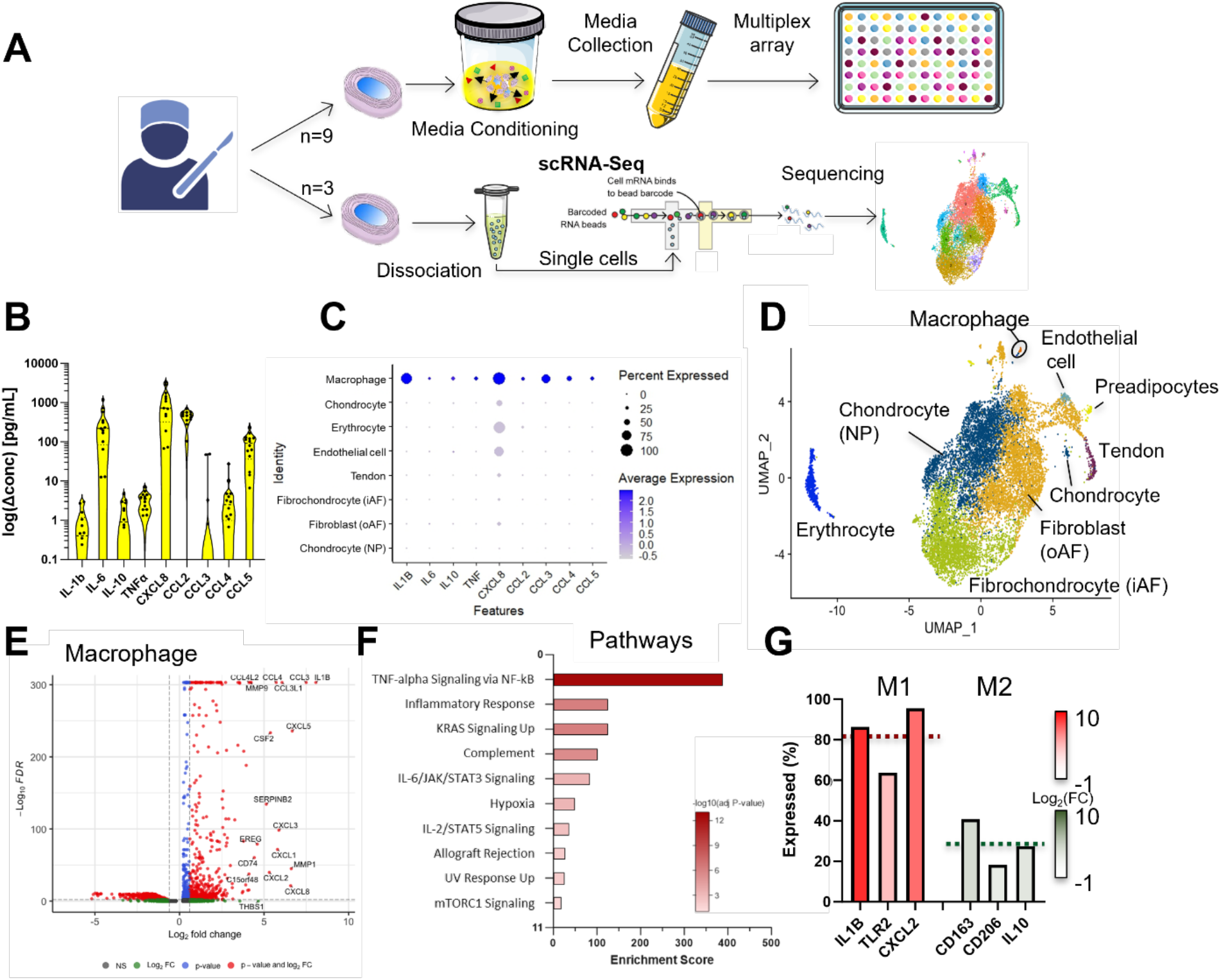
Painful human IVDD tissue contains high concentrations chemokines and cytokines, dominantly expressed from a small population of M1 macrophages highly enriched for TNF-alpha signaling. (A) Specimen from subjects were collected and either used for CM generation and characterization (top) or for cell dissociation and scRNA-seq analysis (bottom). (B) IVDD CM contains a complex array of pro- and anti-inflammatory cytokines and chemokines. (C) IVDD cytokines are dominantly expressed by macrophage sub-population of IVD cells with surprisingly little expression in other IVDD cell populations. (D) UMAP of IVDD cells annotated with UniCell Deconvolve identified 7 cell types including a small population of macrophages. Since relatively few IVD studies are included in the UniCell Deconvolve training set, we parenthetically added IVD-specific annotations to the UMAP. (E) Volcano plot shows DEGs with pro-inflammatory genes CCL3, CCL4 and IL1B as most significantly upregulated genes, which contribute to (F) pathways analysis revealing TNF-alpha signaling as the most significant and highest enriched pathway of this population. (G) M1/M2 macrophage polarization quantification showed a 3-fold higher amount of M1 macrophages within the IVD.

Cell cluster 13 was identified as a small population of macrophages using UniCell and we manually confirmed this annotation as macrophages by evaluating expression of markers including *CD68, CXCL16, C1QB* and *CD14* (Fig. S2A). While these macrophages represent only 0.16% of the cells of the specimens (Fig. S2B), they are highlighted because they expressed the genes translating to the cytokines present in the IVDD CM at high levels (Fig. 1C, Fig. 1D). Specifically, the macrophage cluster contained 217 significantly upregulated genes including *IL1B, CCL3* (*MIP1α*), *CCL3L1* and *CCL4* (*MIP1β*), and emphasized the robustness of its pro-inflammatory roles compared to all other cell clusters (Fig. 1E), and this population did not express IVD cell phenotypic markers. Pathway analysis showed macrophage expression pattern was significantly enriched for “TNFα Signaling via NFκB” (Score= 385.8, adj. p-value= 2.27x10^-13^), “Inflammatory Response” (Score= 123.6, adj. p-value= 5.89x10^-7^) and “Complement” (Score= 100.3, adj. p-value= 3.22x10^-6^) (Fig. 1F). M1-macrophage markers were 2.5-fold greater (average of 82%, red dotted line) than M2-macrophage markers (average of 29%, green dotted line) suggesting macrophages were in an M1 pro-inflammatory phenotype with little conversion to M2 reparative phenotype (Fig. 1G, Fig. S2C). Macrophages expressed the cytokines found in CM at high levels suggesting a role contributing to the CM profiles. Meanwhile, cytokines in CM were expressed by IVD cells at levels that were considerably lower than by the macrophages. We conclude IVDD tissue from back pain subjects contains a complex ‘cocktail’ of pro-inflammatory cytokines and chemokines dominantly expressed by a very small population of macrophages enriched for TNFα signaling.

### CM inhibited hAF cell metabolism and produced a muted inflammatory response

We next used CM to challenge hAF cells in the development of an *in vitro* model system representing the complex IVDD cytokine microenvironment (Fig. 2). Use of the CM cytokine ‘cocktail’ was significant since most IVD cell culture models challenge cells with a single cytokine (most commonly TNFα or IL-1β). The 2D hAF cell culture model system maintains the elongated phenotype of native AF cells and simplifies cellular solute transport conditions. CM was applied in a 2-part study on dosing and timing, and compared with Basal media. CM was generated from specimens collected and pooled from 9 subjects (Table S1). The dosing study applied increasing CM concentrations (Basal, 25%, 50% and 100% CM) to hAF cells cultured for 24 hours in Basal media (Fig. 2A). CM concentration decreased the number of hAF cells visible in culture (Fig. 2B) and significantly decreased hAF cell metabolic activity (Fig. 2C). While increasing CM dose reduces relative nutrients available, we confirmed glucose concentration in all CM conditions was several folds greater than required for hAF cells under these culture conditions (46) to confirm results were related to cytokine concentration changes.

**Fig. 2.**
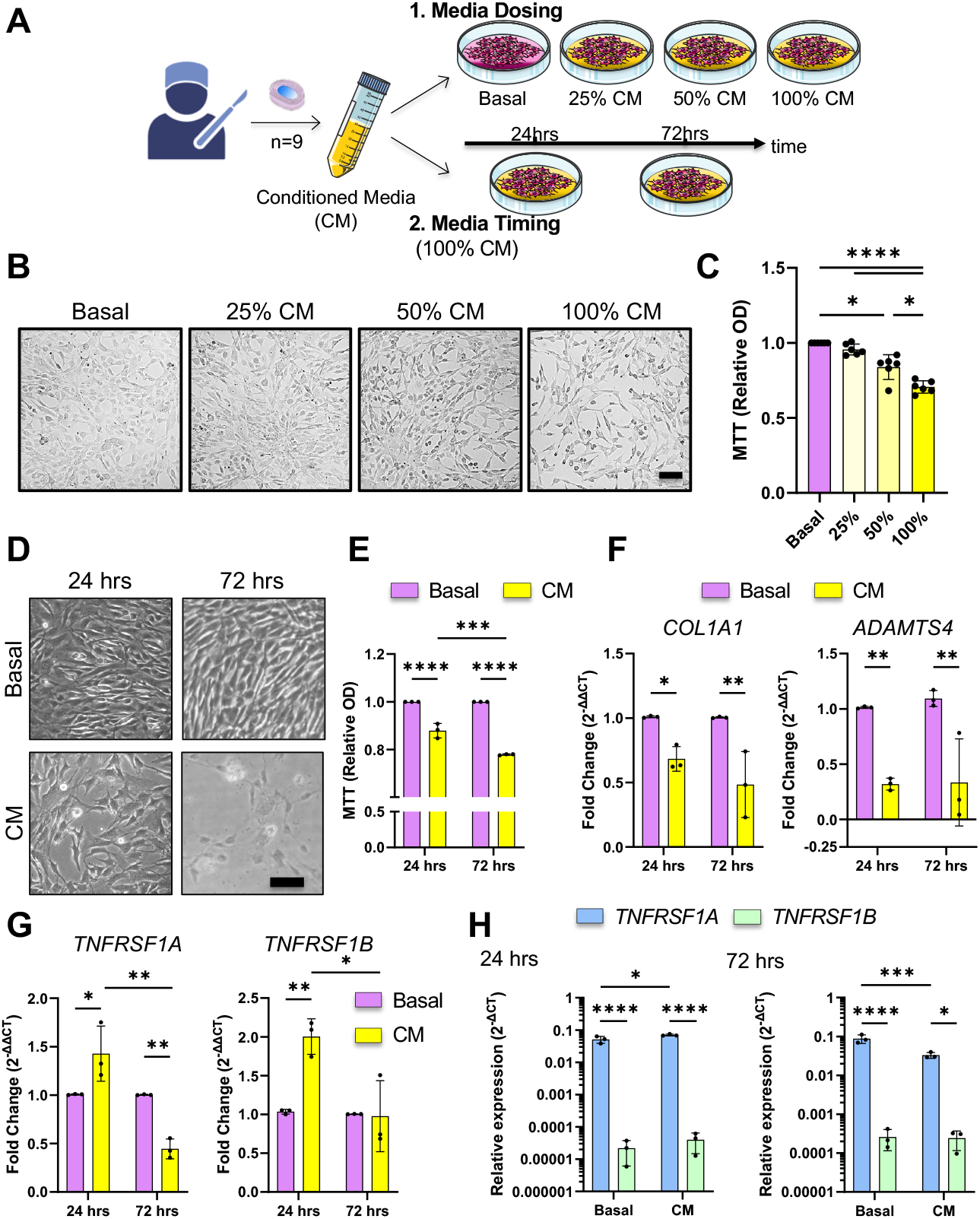
Human AF cell IVDD model system shows CM reduces MTT metabolic activity and modulates TNFR1 and TNFR2 responses. (A) CM generated from 9 subjects were first applied in different dosages (0-100%) for 24 hours (“Media Dosing”), followed by culture for 24 hours and 72 hours respectively (“Media Timing”) (B) Cell growth area decreases with increasing concentration of CM, which is reflected in (C) where cell proliferation significantly decreases with increasing CM, indicated by MTT assay. (D) Culturing hAF cells for 72 hours in CM has a detrimental effect on cell growth and morphology compared to Basal and 24 hours CM exposure, which is also seen in (E) with significantly lower relative OD, indicating lower cell growth, and (F) reduced *COL1A1* and *ADAMTS4* expression, and limited inflammatory response (Fig. S3). (G) hAF cell culture in CM for 24 hours significantly increases gene expression *TNFRSF1A* (gene for TNFR1) and *TNFRSF1B* (gene for TNFR2) compared to Basal and 72 hours culture. (H) Relative gene expression normalized to *GAPDH* of *TNFRSF1A* and *TNFRSP1B* after 24 and 72 hours (2^-ΔCT^). *, **, *** and **** represent significant differences between groups with p< 0.05, 0.01, 0.001 and 0.0001 respectively. Scale bar = 100 µm.

We next performed a timing study with 100% CM (Fig. 2A) and showed CM treatment again reduced cell numbers and MTT at 24-hours and further decreased cell numbers with more granulated cell morphology with elongated stress fibers (Fig. 2D), and reduced MTT at 72 hours (Fig. 2E). *COL1A1* and *ADAMTS4* were also significantly decreased with CM at both time points (Fig. 2F), although CM exposure did not result in significantly altered pro-inflammatory cytokines (Fig. S3C). *TNFRSF1A* and *TNFRSF1B* (genes for TNFR1 and TNFR2, respectively) were significantly increased with CM at 24 hours but then significantly decreased at 72 hours (Fig. 2G). Even though *TNFRSF1A* and *TNFRSF1B* levels were both responsive to CM treatment, we show *TNFRSF1B* was expressed at extremely low levels 100-1000x times below *TNFRSF1A* expression levels (Fig. 3H). Overall, we conclude that CM causes hAF cells to have reduced metabolic rate, and a limited pro-inflammatory response. We also identified that CM altered TNFR1 and TNFR2 signaling, yet, hAF cells expressed *TNFRSF1B* at levels orders of magnitudes lower than *TNFRSF1A*.

**Fig. 3.**
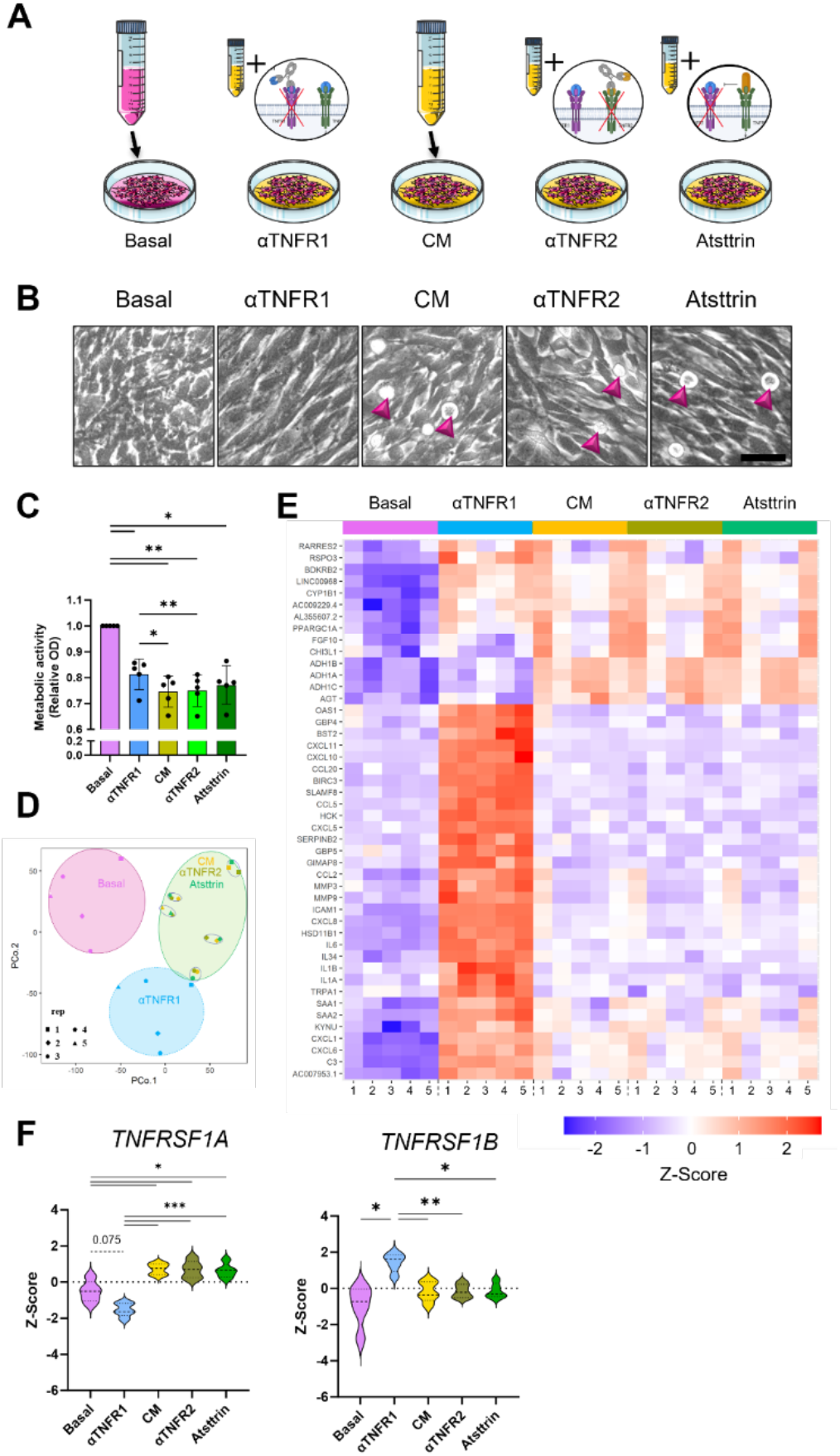
TNFR1 blocking of hAF cells with IVDD CM partially rescues metabolic rate and expresses a pro-inflammatory transcriptome while TNFR2 blocking and Atsttrin had no effects. (A) Cells were either cultured for 24 hours in Basal media, CM or CM supplemented with αTNFR1, TNFR2 or Atsttrin respectively. (B) hAF cells in CM, αTNFR2 and Atsttrin show greater number of floating dead cells (arrowheads), which is consistent with (C) reduced cell growth/viability in CM, αTNFR2 and Atsttrin compared to basal, which was partly recovered in αTNFR1. (D) Distinct transcriptomic difference between Basal, αTNFR1 and all other groups are apparent in PCA plot, and (E) heatmap, which shows a clear change in top DRG patterns with many pro-inflammatory genes among the top 50 DEGs. (F) Gene expression of TNFR1 is significantly downregulated upon TNFR1 blocking, whereas TNFR2 is upregulated. *, ** and *** represent significant differences between groups with p< 0.05, 0.01 and 0.001 respectively. Scale bar = 50 µm.

### TNFR1 blocking enhanced hAF cell metabolic and pro-inflammatory response while TNFR2 blocking and Atsttrin had no effects

Since CM altered expression levels of *TNFRSF1A* and *TNFRSF1B* in hAF cells, we next evaluated their functional roles on hAF cells challenged by CM using blocking antibodies αTNFR1 and αTNFR2, and using TNFR2 activator Atsttrin with bulk RNA-Seq analyses. CM was supplemented with inhibiting antibodies at 10 µg/mL and Atsttrin at 200 ng/mL, and hAF cells were cultured for 24 hours (Table 1, Fig. 3A). Blocking TNFR1 partially rescued hAF cells from CM with improved cell morphology (Fig. 3B) and significantly increased MTT (p <0.05, Fig. 3C) compared to CM (p<0.05) and αTNFR2 (p<0.01), although αTNFR1 was significantly reduced compared to Basal (p<0.05, Fig. 3C). While MTT values were only partially rescued for αTNFR1, it was sufficient to promote a substantially different transcriptome with robust pro-inflammatory transcriptomic profile (Fig. 3D & 3E). Specifically, Bulk RNA-seq with principal component analysis (PCA) on transcriptome showed three distinct clusters (Basal, αTNFR1 and the three remaining groups – CM, αTNFR2, Atsttrin) (Fig. 3D). The αTNFR1 robustly increase in pro-inflammatory cytokines expression shown on the top 50 DEGs on the heatmap (Fig. 3E). *TNFRSF1A* expression was significantly upregulated in CM, αTNFR2 and Atsttrin compared to Basal, and significantly inhibited for αTNFR1, validating the importance of TNFR1 signaling under CM conditions and successful blocking by αTNFR1 (Fig. 3F). *TNFRSF1B* expression was not affected by CM, αTNFR2, or Atsttrin suggesting little TNFR2 signaling under the complex cytokine challenge of CM.

**Table 1.**
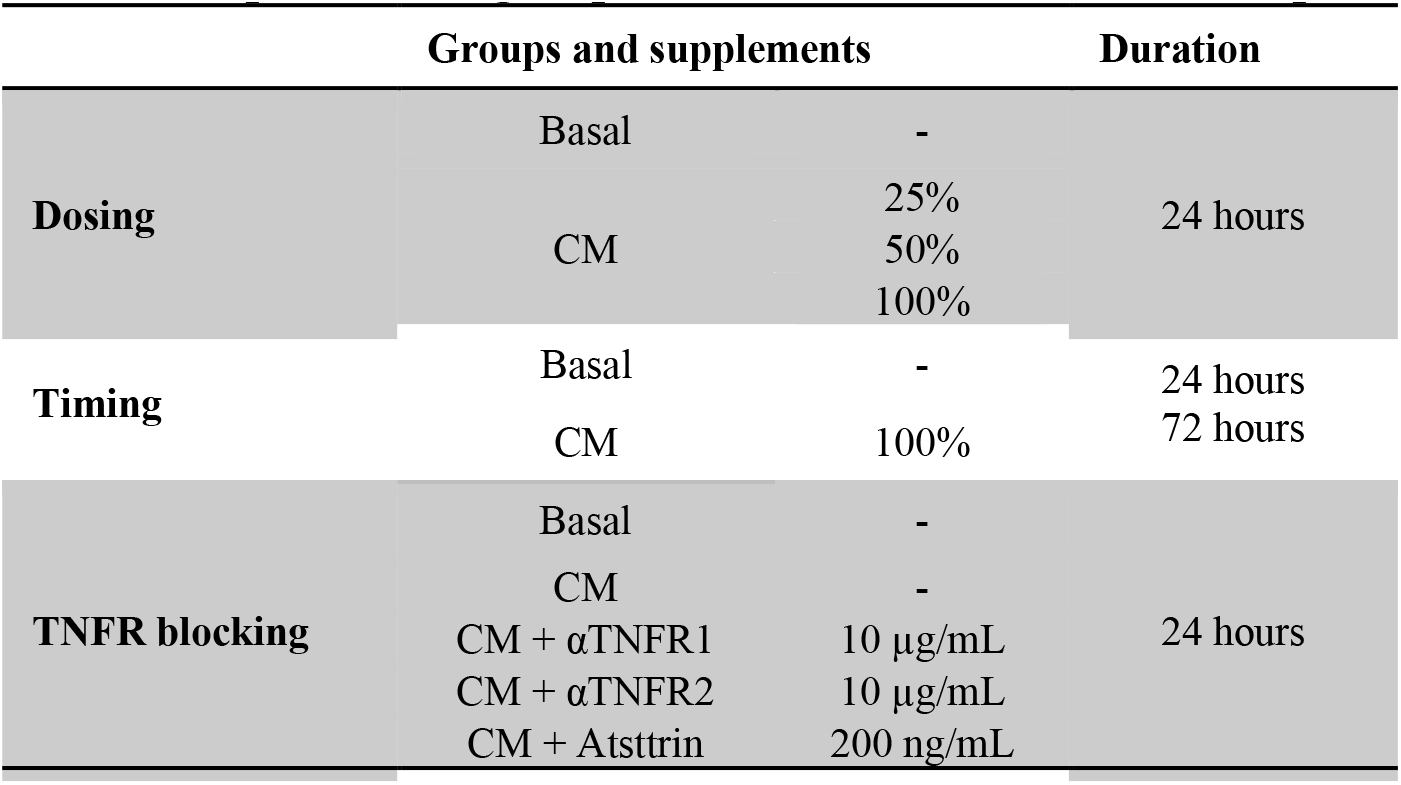
Experimental groups for *in vitro* hAF cell culture experiments

CM, αTNFR2 and Atsttrin groups had nearly identical responses for all assays, suggesting little TNFR2 signaling. Specifically, CM, αTNFR2 and Atsttrin exhibited more floating cells compared to Basal and αTNFR1 (pink arrows, Fig. 3B) and significantly lower MTT relative optical density compared to Basal (Fig. 3C). The majority of the variability on the green cluster on PCA was from donor differences with very little variability from treatment group (CM, αTNFR2 or Atsttrin) (Fig. 3D). The heatmap of the top 50 differentially expressed genes (DEGs) similarly showed distinct patterns for Basal, αTNFR1 and CM groups and near identical patterns for CM, αTNFR2 and Atsttrin groups (Fig. 3E). While the Z-score for *TNFRSF1B* was significantly increased with αTNFR1 inhibition (Fig. 3F), we believe TNFR2 signaling may not have biological impact in IVD since it was not reduced by αTNFR2, since CM, αTNFR2 and Atsttrin had nearly identical results across assays, and because qPCR showed *TNFRSF1B* expression levels were 100- 1000x lower than *TNFRSF1A* (Fig. 2H). Overall, we have 3 key findings. First, CM significantly inhibited metabolic rate of hAF cells. Second, TNFR1 inhibition partially restored hAF cell metabolic rates sufficiently to enable a robust pro-inflammatory response. Third, results suggest little functional role for TNFR2 signaling in hAF cells exposed to CM. The lack of responsiveness of TNFR2 inhibition adds to our data on lack of TNFR2 expression in IVD cells in supporting our hypothesis that IVD cells have limited TNFR2 signaling.

### CM inhibited hAF DNA replication while αTNFR1 shifted hAF cell pro-inflammatory response to predict enhanced immune cell recruitment

Basal, αTNFR1 and CM were analyzed further with gene set enrichment analysis (GSEA) and network analysis with ingenuity pathway analyses (IPA) because of the large transcriptional differences and to discern how TNFR1 modulation shifted the response of hAF cells to the CM challenge. Meanwhile αTNFR2 and Atsttrin were not further analyzed since they had negligible transcriptomic differences from CM. CM significantly downregulated 509 DEGs and significantly upregulated 762 DEGs compared to Basal (Fig. 4A). The most significantly downregulated DEGs were of the minichromosome maintenance gene family (*MCM* genes), which play roles in DNA replication (47). GSEA further showed CM had the highest scores for depleting biological processes related to Cell cycle and DNA replication while inflammatory responses were most significantly enriched (Fig. 4B). The αTNFR1 treatment compared to CM significantly downregulated 490 DEGs, and significantly upregulated 545 DEGs (Fig. 4C). GSEA predicted these DEGs to be enriched for IL-27 mediated signaling (Fig. 4D), which has multiple effects in modulating inflammatory responses (48). Additional enriched biological processes were related to Regulation of Neuroinflammatory Response, and Response to Cytokine, demonstrating a shift in the inflammatory pathways activated by CM (Fig. 4D). Processes related to Presynaptic Endocytosis and Regulation of Calcium Ion Dependent Exocytosis were significantly depleted, suggesting TNFR1 blocking could influence communication with the peripheral nervous system as compared to CM. The αTNFR1 group (which also contained CM) compared to Basal significantly downregulated 448 DEGs and significantly 1072 DEGs with *NFKB2*, *TNFAIP3* and *CCL5* among the highest fold change (FC) DEGs (Fig. 4E). The αTNFR1 treated CM condition, therefore enriched biological processes related to cellular response to type 1 interferon, cytokine-mediated signaling pathways, and inflammatory response compared to Basal and depleted processes related to axonogenesis and axon guidance (Fig. 4F). Network analysis using QIAGEN IPA predicted that CM would activate hAF cell senescence and inhibit proliferation compared to Basal (Fig. 4G). Meanwhile network analyses predicted that hAF cells treated with αTNFR1 would enhance cell activity and processes related to immune cell recruitment compared to CM (Fig. 4H). Further, αTNFR1 treated CM predicted inhibition of SOCS1, a suppressor of cytokine signaling, compared to Basal, to promote the robust pro-inflammatory response observed. Overall, CM depletes DNA replication and enriches inflammation. While, αTNFR1 enabled enhanced the hAF cell pro-inflammatory response to CM and predicted immune cell recruitment.

**Fig. 4.**
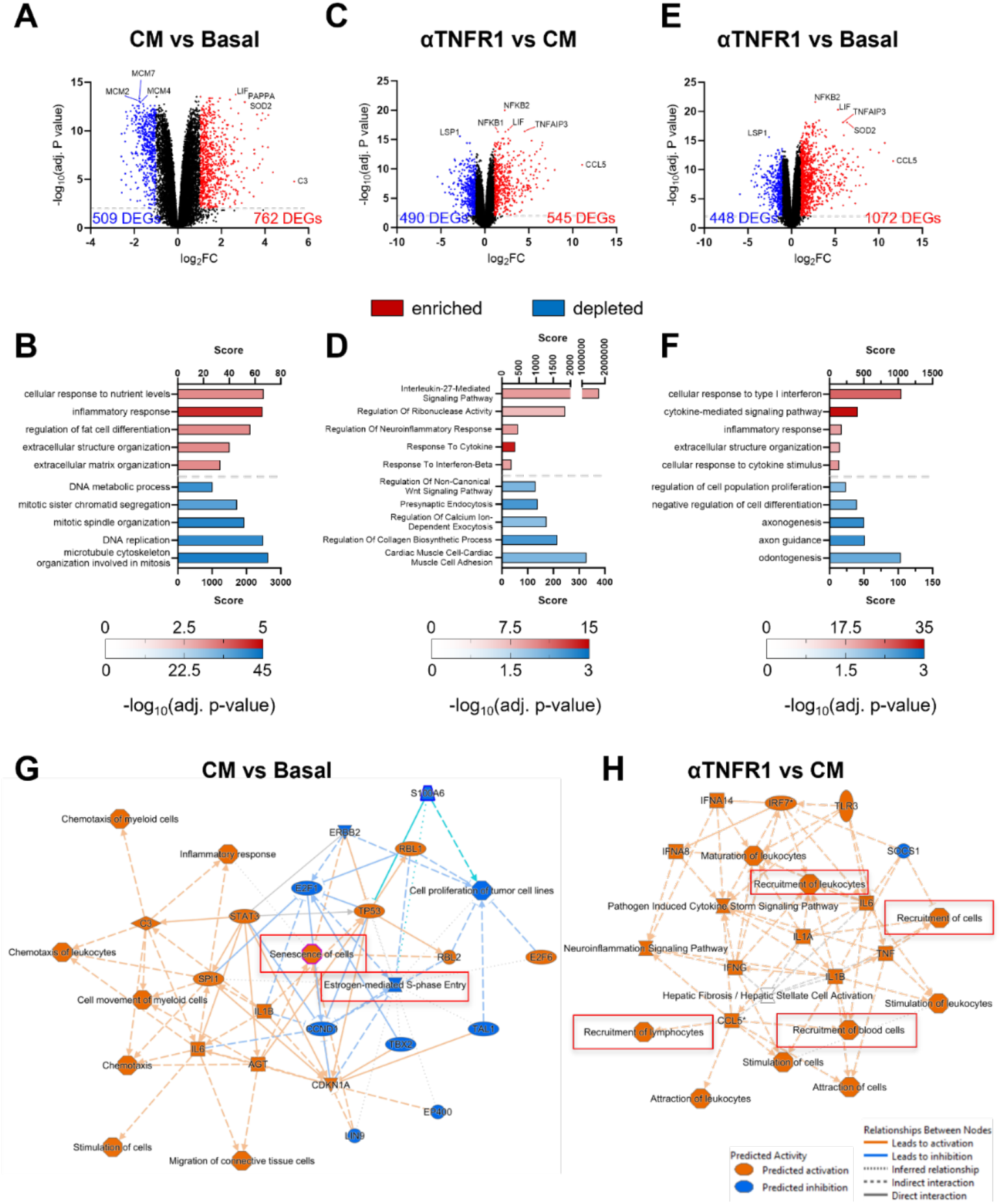
CM reduces mitotic responses while TNFR1 inhibition promotes processes for immune cell recruitment. (A) 509 of 1271 DEGs in CM were significantly downregulated compared to Basal. (B) Biological processes depleted upon CM were related to DNA replication and mitosis. (C) Comparing αTNFR1 to CM, 454/1035 DEGs were significantly upregulated. (D) Biological processes modulated by αTNFR1were related to an enrichment of IL-27 signaling and RNase activity. (E) Volcano plot comparing αTNFR1 to Basal significantly upregulated 1072/1520 DEGs. (D) Hereby, biological processes related to cytokine signaling and inflammatory responses were enriched, while axonogenesis and axon guidance was depleted. (G) Qiagen IPA analysis of CM vs Basal predicted an activation of senescence while proliferation was inhibited. (H) Inhibition of TNFR1 activates processes related to immune-cell recruitment in IPA analysis.

### CM inhibited cell cycle and promoted senescence that was partially restored with αTNFR1

The reduced metabolic rates of hAF cells with increased CM concentration and reductions with αTNFR1 motivated a closer investigation of the cell proliferation, apoptosis, and cell cycle for Basal, αTNFR1, and CM groups (Fig. 5A). We considered cell cycle genes transcription patterns relevant across multiple cellular models across different biological contexts (49, 50), and determined that CM significantly downregulated hAF genes related to each phase of the cell cycle compared to Basal (Fig. 5B). Furthermore, CM caused hAF cells to be express G0 markers indicative of quiescence (i.e., downregulation of *FOXM1, BIRC5* and *H2AFZ* and upregulation of *PFDN5* and *RPS14*) (50) relative to Basal. Blocking TNFR1 led to a pattern of cell cycle markers between CM and Basal, suggesting a partial cell cycle recovery. *MKI67*, the gene coding for the proliferation marker KI67, was significantly decreased for hAF cells in CM (p<0.0001), while αTNFR1 was significantly different from CM and Basal (Fig. 5C). The senescence marker SA-β- Gal (Fig. 5D) was significantly increased in hAF cells in CM compared to Basal, and compared to αTNFR1 (Fig. 5E), again suggesting a partial recovery by αTNFR1. The apoptosis marker Caspase-3 (Casp3) (Fig. 5F) showed no difference in co-localization of cleaved Casp3 and DAPI using the Pearson’s correlation coefficient (PCC) or in fluorescent intensity within the nuclei between groups (Fig. 5G, Fig. 5H), suggesting no changes in apoptosis. Together, transcriptomic and protein assay results indicated that CM downregulated hAF cell cycle markers and increased senescence and not apoptosis, while αTNFR1 partially protected and restored cell cycle from senescence. While the senescence and cell cycle recovery was only partial with αTNFR1, the expression profile (Fig. 3, Fig. 4) showed it was sufficient to enable hAF cells to mount a robust pro-inflammatory response.

**Fig. 5.**
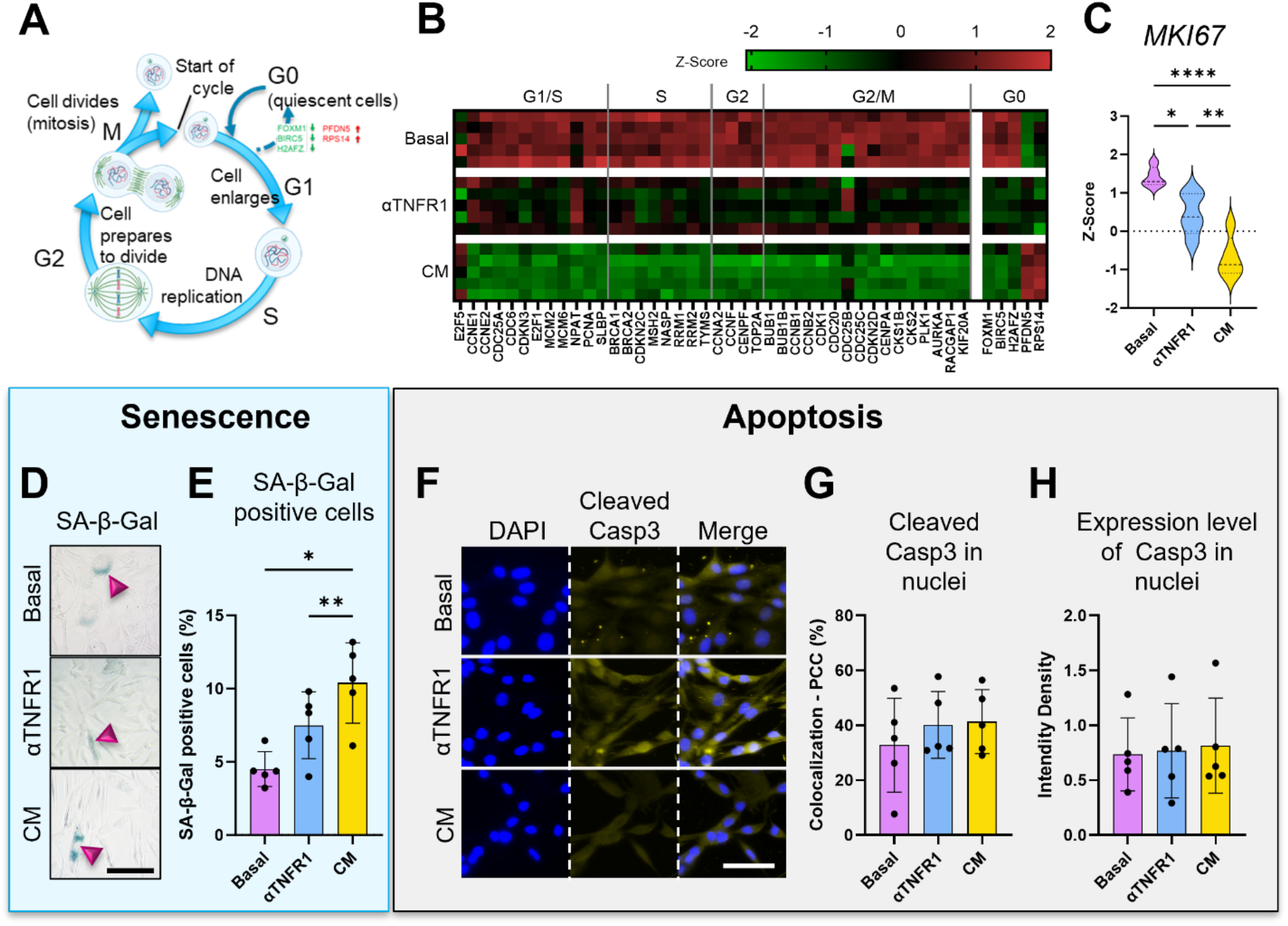
IVDD CM inhibited active hAF cell mitosis and promoted senescence that was partially prevented with TNFR1 inhibition. (A) Schematic of different stages of the cell cycle. (B) Heatmaps using transcripts selected to study cell cycle in multiple cellular models and different biological contexts (49, 50) showed Basal activated hAF cell cycle while CM caused expression of G0 phase transcripts. (C) MKI67 proliferation marker was significantly decreased in CM compared to Basal and αTNFR1. (D) SA-β-Gal staining of hAF cells was (E) significantly higher in CM compared to Basal and αTNFR1. (F) Staining for cleaved Casp3 showed (G) no difference in co-localization of cleaved Casp3 and DAPI and (H) no difference nuclear cleaved Casp3 intensity density. *, ** and **** represented p< 0.05, 0.01 and 0.0001, respectively. Scale bar = 50 µm.

### TNFR1 increases with IVDD while TNFR2 is expressed only in macrophages

We further analyzed roles of TNFR1 and TNFR2 in IVDs using immunohistochemistry (IHC) on human autopsy samples and scRNA-seq on human IVDD samples. Human IVDs of various degeneration grades (n=14, Rutgers grade 1.3-12) were stained for TNFR1 and TNFR2 respectively and the outer AF (oAF) analyzed using HALO Multiplex module (Fig. 6A). TNFR1 was found in the cell membrane of oAF cells as expected, and the % positive cells for TNFR1 significantly correlated with IVDD Grade (Fig. 6B, Fig. 6C). This significant correlation between TNFR1 protein expression and Rutgers IVDD grade was also found in the inner AF and NP (Fig. S5). TNFR2 only showed faint staining within and around the nuclei of oAF cells suggesting little presence of TNFR2 (Fig. 6B). Furthermore, the % TNFR2 positivity of these faintly stained cells were not affected by IVDD grade in the oAF (Fig. 6D), or inner AF and NP regions (Fig. S5). We used scRNA-seq of human IVDD subjects to further investigate which human IVD cells expressed the genes for TNFR1 and TNFR2. *TNFRSF1A* was expressed in nearly all cells in the scRNA-seq analyses and had particularly high expression levels in AF and NP cells as well as erythrocytes (Fig. 6E, Fig. 6F) supporting its important role in IVD response to IVDD CM conditions. In contrast, *TNFRSF1B* was at extremely low levels in all IVD cell populations (Fig. 6E, Fig. 6F), in line with the extremely low expression levels in hAF cells on qPCR, and faint TNFR2 staining. Interestingly, the small macrophage population expressed *TNFRSF1B* at very high levels and *TNFRSF1A* at negligible levels. Results therefore support our hypotheses that TNFR2 signaling is very low in all IVD cells and further emphasize the importance of TNFR1 signaling in all IVD cells with TNFR1 signaling increasing with IVDD grade. The scRNA-seq also shows importance of TNFR2 signaling in immune cell populations.

**Fig. 6.**
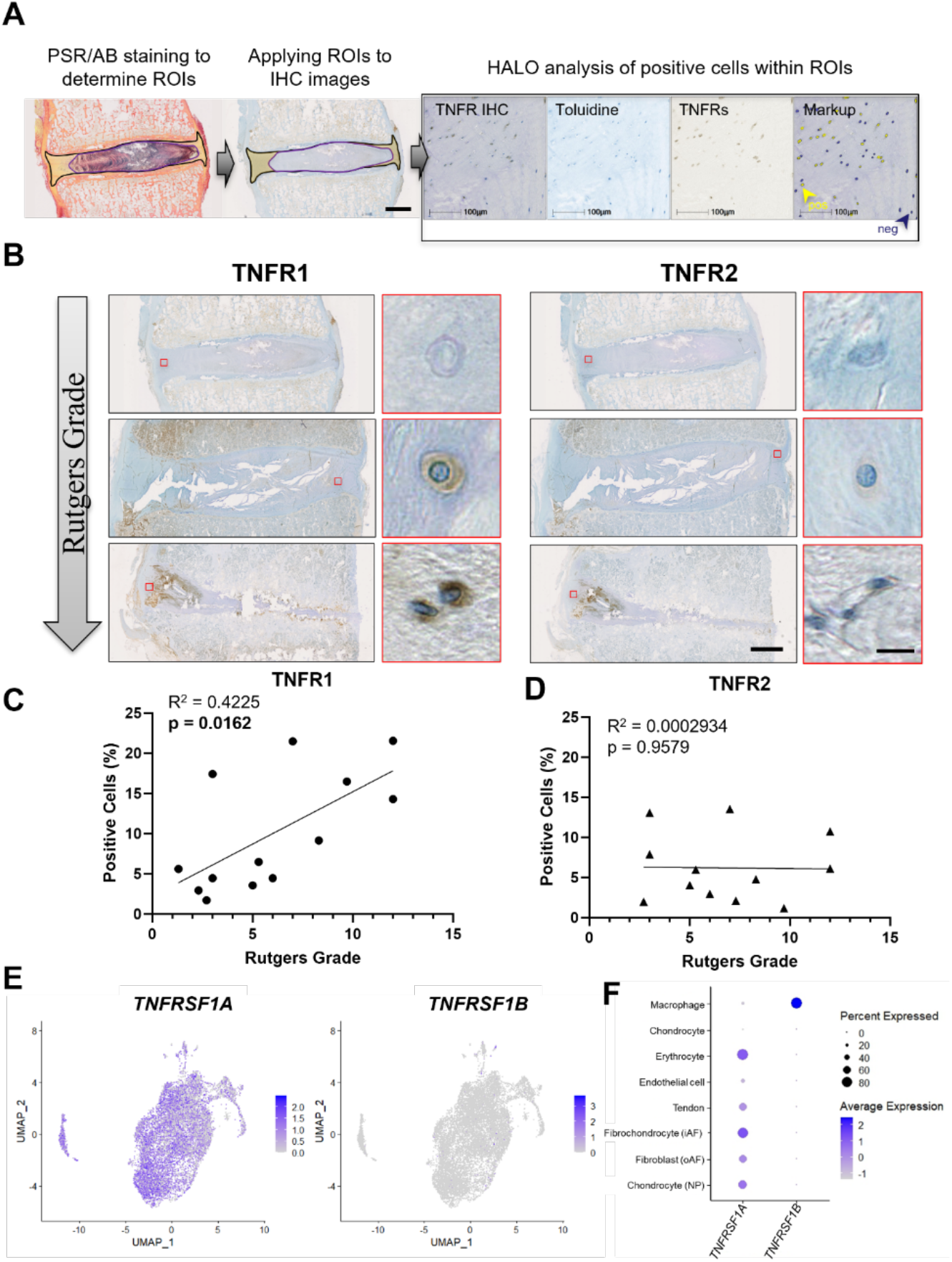
TNFR1 expression was expressed in all IVD cells and correlated with IVD degeneration grade while TNFR2 was only expressed in macrophages and not IVD cells. (A) Analysis of IHC was performed using HALO Multiplex module, whereby the oAF region was determined using PSR/AB staining and the ROI overlayed onto TNFR1/2 IHC imaged for follow up AI-based analysis. (B) Intensity and quantity of TNFR1 increases with Rutgers degeneration grade, while TNFR2 shows faint staining across degeneration. (C) Correlation between TNFR1 positive cells and Rutgers grade is significant (p=0.0162), while (D) there is no correlation with TNFR2 positive cells and Rutgers grade. (E) UMAP of herniated IVD specimen showed TNFR1 expression across cell populations and little TNFR2 expression. (F) TNFR2 is expressed exclusively in macrophage-derived cell population. Scale bar (full IVD) = 5 mm, Scale bar (AF cell) = 10 µm.

## DISCUSSION

There is a major need to develop improved therapeutic targets to manage the chronic pro-inflammatory conditions in painful IVDD, and to develop strategies for IVD cell immunomodulation to slow IVDD progression or enhance IVD healing. Here, we use multiple human model systems to understand roles of TNFR1 and TNFR2 in modulating the complex cytokine and chemokine environment of IVDD. We first show that IVDD derived CM contains elevated levels of at least 32 cytokines and chemokines highlighting the complex pro-inflammatory environment with known roles recruiting inflammatory cells. It was notable that these chemokines were dominantly expressed by a small population of macrophages while native IVD cells (present in substantially larger numbers) expressed these pro-inflammatory factors at much lower levels. CM treatment on hAF cells had the largest effects reducing metabolic rate, and promoting cell cycle arrest and senescence, and promoted more muted transcriptional expression patterns. TNFR1 signaling was strongly involved in this muted metabolic and inflammatory response since results showed TNFR1 was expressed by all IVD cells and increased with IVDD grade, and since blocking TNFR1 with a CM challenge rescued hAF cell metabolic rate and senescence sufficiently to mount a robust pro-inflammatory response. On the other hand, results supported the hypothesis that TNFR2 signaling was very limited in IVD cells since blocking and Atsttrin had no effects on hAF treated with CM treatment, and since extremely low levels of TNFR2 were measured in IVD cells with qPCR, bulk RNA-seq, scRNA-seq and IHC staining. Meanwhile macrophages expressed TNFR2 at high levels, and αTNFR1 treatment predicted enhanced inflammatory cell recruitment suggesting the immunomodulation of TNFR1 and TNFR2 signaling in the IVD may be possible with enhanced interactions of IVD and immune cells.

We showed that IVDD tissue from herniation subjects produced high concentrations of several pro-inflammatory cytokines and chemokines that had their largest effects limiting hAF cell metabolic and inflammatory responses. The complex combination of cytokines and chemokines present in IVDD CM are consistent with literature showing a variety of cytokines increase with severity of symptomatic IVD herniation, and that elevated IL-6 plays an important role in painful conditions (11, 51). CM was used to establish and characterize an *in vitro* hAF cell IVDD model capable of promoting and maintaining an IVDD phenotype. The major effects of CM on hAF cells were reduced IVD cell metabolic rates, arrested cell cycle, senescence, and a relatively muted transcriptional response to the complex cytokine and chemokine mixture in CM conditions. Inhibiting TNFR1 enabled hAF cells to sufficiently recover from CM conditions to mount a robust pro-inflammatory response suggesting recovery may be possible under certain conditions. The IVD extracellular matrix is capable of binding cytokines such as TNFα and IL-1β through fibronectin (52), and these cytokines sequestered within the IVD matrix may continue stimulating an IVDD response on affected cells. Individual cytokines can induce premature senescence in IVD cells (17, 53) and this study further shows complex cytokines in CM can arrest hAF cell cycle, induce senescence, and limit IVD cell inflammatory responses independent of nutrient load. The harsh IVDD chemical microenvironment with limited nutrients is often considered a key factor in the ‘frustrated’ IVD healing response that stimulates the production of chronic pro-inflammatory conditions and results in increased senescence observed with IVDD (54–60). However, IVD cells are adapted to the low oxygen and glucose microenvironment of the IVD and limited nutrients do not explain the reduced hAF cell metabolism from CM challenge in this cell culture study since there were ample nutrients present in CM conditions of all concentrations. We used Basal as control because healthy IVD controls were not available (i.e., surgical samples are degenerated, scoliosis is a different disease process, we do not have access to organ donor samples, and autopsy samples are treated differently and not comparable), and we expect healthy IVD CM would produce effects between the Basal and IVDD CM conditions reported in this study. This study supports the literature showing IVDD cytokines and chemokines play roles inhibiting IVD cell cycle, promoting senescence (17), and inducing a muted transcriptional response in IVDD.

IVD cell immunomodulation may be important to arrest IVDD and promote IVD healing, and this study strongly showed a role for TNFR1 since blocking TNFR1 enable hAF cells to mount a robust pro-inflammatory response to the complex CM mixture, even though cell metabolic rate and senescence were only partially rescued. Wang et al., showed a mechanistic role of TNFR1 in IVD healing with significantly improved IVD degeneration score in TNFR1^-/-^ mice (20). The strong TNFR1 immunostaining and increased % positivity with advancing human IVDD grade in this study is similar to prior studies on TNFR1 IHC (31, 61, 62). Translationally, TNFR1-CRISPRi modulation in a rat *in vivo* model improved IVD degeneration and pain response (63). Reduced pain-like behaviors with TNFR1 inhibition are consistent with the depletion of axonogenesis and presynaptic endocytosis identified in this study when blocking the TNFR1 pathway, and suggests TNFR1 may play roles in pain and IVDD. The more robust pro-inflammatory response in this study with TNFR1 inhibition may be beneficial since recent studies suggest acute inflammation is important to promote healing and inhibit chronic pain (64). Further, regenerative healing in tendons requires a more robust and short-term immune response compared to inferior adult healing (65, 66). We believe this study and growing literature suggest that IVD cell immunomodulation with a more acute pro-inflammatory response may be important in shifting IVDD conditions to a more reparative state, and this study builds on the literature suggesting TNFR1 modulation as an important therapeutic target.

The limited TNFR2 signaling in IVD cells found in this study was a strong finding supported by qPCR, bulk RNA-seq, scRNA-seq, IHC methods and blocking studies. Lack of TNFR2 signaling in IVD cells has potential for a shift in our understanding of IVD pathophysiology, suggesting IVD cells have limited capacity to mount a repair/regeneration response regardless of their microenvironmental conditions. However, there is a significant body of literature on TNFR2 signaling in the IVD that lacks clarity and needs to be contextualized with this study. For example, TNFR2 IHC was expressed in up to ∼ 15% of IVD cells in this study, but the immunostaining was very faint and had no changes in IVD cells with advancing IVDD. Even though review articles suggest TNFR1 and TNFR2 are comparably expressed in the IVD (9, 10), the papers cited are more closely aligned with our findings suggesting limited role for TNFR2. Specifically, TNFR2 was not detectable in AF and NP samples from controls or at very low expression levels and was not increased with IVDD (31, 61, 62). However, TNFR2 levels were increased in human IVDs near defects and in IVDs with immune cells infiltrates (61, 62), suggesting increased TNFR2 signaling in IVDs involves immune cells. The scRNA-seq in this study results support this concept showing *TNFRSF1B* in inflammatory cells but not in IVD cells, and this is supported by our qPCR results showing extremely low *TNFRSF1B* levels in hAF cells. Furthermore, the PanglaoDB scRNA-seq database with over 5M mouse and human cells sequenced shows *TNFRSF1B* is expressed at detectable levels in few cell types, predominantly immune cells including dendritic cells, macrophages and T cells (67). We believe the lack of effects of αTNFR2 and Atsttrin in hAF challenged with CM in this study is best understood if hAF cells indeed have negligible TNFR2 signaling since αTNFR2 had no effects and since Wang et al., showed that Atsttrin had no effects on TNFR2^-/-^ mice to indicate that Atsttrin is TNFR2-dependent (20). However, TNFR2 signaling is important in IVD degeneration and healing since IVD degeneration increased in TNFR2^-/-^ mice and Atsttrin suppressed TNFα-mediated inflammation in IVD tissue and organ culture models (20, 37). The model systems of these studies showing TNFR2 efficacy in IVDD involve immune cells that are capable of robust TNFR2 signaling. Therefore, our results and the literature all point to a very limited role of TNFR2 in IVD cells. However, TNFR2 signaling appears to be important in IVDD progression and most likely involves TNFR2 signaling of macrophages and possibly other immune cells.

The scRNA-seq analyses identified that the cytokines detected in the multiplex array of IVDD CM were dominantly expressed by a very small population of macrophages and only expressed by native IVD cells at drastically lower levels. Despite substantially lower expression levels in IVD cells, their contribution to the CM cytokines cannot be excluded due to their larger cell numbers in the tissue. The small population of macrophages present were in a predominantly M1 phenotype and also expressed markers (i.e*., ACP5* and *CTSK*, the genes coding for the proteins TRAP and CATHEPSIN K), suggesting their potential osteoclastic origin (Fig. S2D, Fig. S2E) (68). The scRNA-seq results in this study are complementary to human IVDs with IHC identifying small numbers of macrophages infiltrating human IVDDs with a predominantly M1 state that does not convert to a reparative M2 polarization state (55). The small numbers of macrophages isolated in human surgical IVDD in this study further suggests there may be too few immune cells infiltrating the IVDs to enable an effective healing response since inflammatory cell numbers and healing responses depend on species, tissue, timing, and age (69). However, human IVD scRNA-Seq studies are mixed in the number of immune cells identified making this a dynamic area of research and some studies have reported greater numbers of inflammatory cells in human IVDs (70–72). The numbers of immune cells identified in human IVDs will vary depending on degenerative state and isolation methods. We also believe differences in relative abundance of immune cells identified across human IVD scRNA-seq studies are likely related to the annotation methods since IVD cells are known to produce many cytokines and exhibit immune-modulatory behaviors (7, 8, 15, 16, 62), and we had a strict definition of macrophages based on UniCell (45) and lack of IVD cell phenotypic marker expression. An integrated data set and improved precision on phenotypic markers in the immune cell populations present in the IVD in healthy and IVDD conditions are needed to better reconcile differences in relative numbers of macrophages identified across IVD scRNA-seq studies. Regardless, immunomodulation of native IVD cells likely offers more promise for translation since IVD cells are present in much greater numbers than immune cells in the IVD. For example, recently developed bioactive tension-activated nanofiber patches can modulate IVD pro-inflammatory responses through the IL-1β pathway (73), and such approaches can be modified to target additional cytokines and receptors to modulate IVD cell response to the complex IVDD CM environment.

In conclusion, results demonstrate distinct roles of TNFR1 and TNFR2 signaling in IVD cell response to IVDD conditions. We show hAF cells subjected to IVDD CM conditions had inhibited metabolic rate, senescence, and a muted inflammatory response. Targeted TNFR1 blocking under the CM challenge partially rescued hAF cell MTT response sufficiently to enable them to mount a more robust inflammatory response to CM challenge, with production of many chemokines capable of recruiting immune cells. Meanwhile, native IVD cells appear to lack the capacity for a TNFR2-driven repair under cytokine challenges and must instead rely on recruiting TNFR2 expressing cells, such as macrophages for repair responses. Results point to future therapeutic strategies for painful IVDD involving immunomodulatory strategies for IVD cells including TNFR1 inhibition to restore IVD cell cycle and IVD cell metabolic rates in response to IVDD conditions, and recruitment or delivery of TNFR2 expressing cells that may offer additional benefit for IVD repair.

## MATERIALS AND METHODS

### Sex as a biological variable

Our study uses human specimen from both sexes for various experiments (outlined below and in supplementary Tables S1 and S3), but it was not considered a biological variable.

### Tissue source

This study was approved by Mount Sinai Institutional Review Board (IRB# STUDY-19- 01394-MOD001) project titled ‘Orthopedic Cell and Tissue Repository’. For media conditioning, nine human IVDs were obtained from patients (6 males, 3 females), 34-65 years old (mean = 50.4). For cell isolation, six IVDs were obtained (4 males, 2 females), 25-71 years old (mean = 45.8). The collected specimen was assessed, and AF and NP were carefully separated for cell isolation. All patients experienced back or neck pain and underwent disc fusion or discectomy surgery. Assessment of the disease state was performed using Pfirrmann grading on Pre-surgery MRI images. Informed consent from patients was given prior to surgery Specimen information is given in Table S1. Human samples for IHC were obtained from an institutional biobank of cadaveric IVDs. This study included 15 lumbar IVDs (8 Male, 7 Female) (Table S3). The morphological degeneration of each sample was quantified using the Rutgers degeneration scale that ranges from grade 1-12, with 12 as the most degenerated (74). Each sample was graded for Rutgers grade by three trained and independent graders who were blinded to the sample age, sex, and to the Pfirrman grade.

### Conditioning of media

Immediately after IVD collection, tissues were cut into smaller pieces, if necessary and washed three times in Phosphate-Buffered Saline (PBS) supplemented with 2% Penicillin / Streptomicin (10,000 U/mL, Gibco) and Primocin (100 µg/mL, Invivogen). Afterwards, the tissue was cultured in 10 mL media (RoosterNourish™-MSC, RoosterBio) per gram tissue and maintained under constant agitation at 37°C and 5% CO2 for 72 hours. CM was collected and frozen as individual samples at -80°C for later analysis. After initial characterization and analysis of statistical differences between samples, media collected from 9 individuals were combined to reduce number of variables in consecutive experiments.

### Media analysis/ Multiplex cytokine array analysis

For quantification of pro-inflammatory cytokines released from surgical samples, media was sampled and analyzed using a Human Cytokine/Chemokine 48-Plex Discovery Assay® Array (catalog no. HD48, Eve Technologies). The 48-Plex Discovery Assay used 100 μL of undiluted cell culture media per run and each run was performed in duplicates. Fluorescence intensity values were derived from the discovery assay and are in direct proportion to the concentration of proteins in the samples.

### Isolation and culture of hAF cells

The hAF cells were isolated from surgical specimens. Briefly, collected AF tissue was finely minced and digested using collagenase type I (400 U/mL, Gibco, USA) for 12 hours under constant agitation at 37°C in serum-free high glucose DMEM containing Penicillin / Streptomycin and Amphotericin B (0.25 µg/mL). Using a 70 µm cell strainer debris was removed from the tissue/cell suspension. Cells were either immediately processed for scRNA-seq analysis or seeded at 5,000 cells/cm^2^ in RoosterNourish™-MSC media (Rooster Bio) under normoxic (21% O2) conditions at 37°C and 5 % CO2 to passage 2-3 to have sufficient cells for CM challenge and blocking experiments. RoosterNourish™-MSC media was selected since it is designed to support high volume expansion of human cells without media changes and therefore had ample nutrients for hAF cell growth under all conditions and obviated the need to adjust nutrient concentrations as we adjusted CM concentration and timing. Specifically, glucose concentration in our media at all CM concentrations ranged 15-40 mM glucose which is several folds greater than the glucose demand for hAF cells in native conditions (46).

When primary cells reach 70-80 % confluency, media was changed according to study design (Table 1). Specifically, for Basal group, media was replaced by fresh RoosterNourish™- MSC media (Rooster Bio), for the dosing experiment, CM was diluted using Basal media and for the TNFR blocking experiment, CM was supplemented with anti-TNFR1 (10 µg/mL, MAB225- 500, R&D Systems), anti-TNFR2 (10 µg/mL, MAB726-500, R&D Systems) or Atsttrin (200 ng/mL, manufactured in-house). Cultures were moved into a hypoxic environment (5% O2) for the duration of the experiment and note that cell growth rates did not result in overlapping cells for the 24-72 hour studies performed.

### scRNAseq sample preparation and analyses

Immediately after isolation, cells of from 3 patients (5 specimens) were further cleaned up using the LeviCell system (Levitas Bio) according to manufacturer’s protocols to remove excess debris. Afterwards, samples were processed according to Chromium 3’ Gene Expression V3.1 Kit (10X Genomics, Pleasanton, CA) using manufacturer’s guidelines. An estimated 5,000 cells were loaded onto each channel with a targeted cell recovery of approximately 3,000 cells/sample. This was followed by sequencing on a S4 NovaSeq chip (Illumina Inc., San Diego, CA). Qubit 3 (Fisher Scientific) and 2100 Bioanalyzer (Agilent Technologies, Santa Clara, CA) were used for quality check of library.

Sequencing data were aligned and quantified using the Cell Ranger Single-Cell Software Suite (version 7.1.0, 10x Genomics) against the reference genome GRCh38. The pipeline grouped and de-duplicated reads confidently mapped to the transcriptome by 10X cellular barcodes and UMIs (Unique Molecular Identifiers), then summarized counts into feature-barcode matrices. The R-based package Seurat (v4.3) was used for further processing and analysis (75). For quality control, standard pre-processing steps were applied to remove cells with <500 mapped genes, <1,000 gene counts, and >20% mitochondrial content. The data were further filtered to removed genes detected in less than 5% of cells per sample. All pre-processed samples were RPCA-integrated using the SCTransform normalization method. The top 4,000 variable features were used to define 20 integration anchors. PCA identified 30 dimensions for unsupervised graph-based clustering via uniform manifold approximation and projection (UMAP) at a 0.5 resolution parameter (76). Genes expressed in >10% of cells in a cluster with an average log2-fold change >0.25 were selected as DEGs using FindMarkers. Cell types identified from the resulting UMAP were manually annotated based on each cluster’s DEG profile and further confirmed using UniCell, an unbiased cell type deconvolution pipeline that leverages a deep learning neural network pre-trained on >28 million annotated single cells from published studies (*43*). Pathway enrichment was performed using Enrichr to determine significantly enriched biological pathways based on upregulated DEGs, and normalized enrichment scores were reported for top gene pathway sets.

### MTT assay

To determine the metabolic activity and respective cell viability, cells were cultured in media, supplemented with 0.5 mg/mL 3-[4,5-dimethylthiazol-2-yl]-2,5-diphenyltetrazolium bromide (MTT, Sigma, USA) for 4 hrs at 37°C. Afterwards, cells were lysed using Dimethyl Sulfoxide to release insoluble formazan, which was quantified at λab = 540 nm with λref >650 nm. Absorbance was normalized to basal group to compare between replicates.

### Gene expression analysis using qRT-PCR

Quantitative reverse transcription polymerase chain reaction (qRT-PCR) was used to quantify the gene expression of hAF cells that were retained during *in vitro* culture of CM model optimization. RNA was extracted from samples using the RNeasy Micro Kit (QIAGEN INC, USA) according to manufacturer instructions and eluted in 15 µL RNase free water. RNA concentration was quantified using a Thermo Scientific NanoDrop™ One UV–Vis spectrophotometer (Thermo Fisher Scientific) followed by synthesis of complementary DNA (cDNA) using the SuperScript® VILO™ MasterMix (Invitrogen, Waltham, MA) according to manufacturer instructions. The PowerUp™ SYBR™ Green Master Mix (Invitrogen) was used with human primers for Glyceraldehyde 3-Phosphate Dehydrogenase (*GAPDH*, Hs.PT.39a.22214836), Collagen Type I Alpha 1 Chain (*COL1A1*, Hs.PT.58.15517795), ADAM Metallopeptidase with Thrombospondin Type 1 Motif 4 (*ADAMTS4*, Hs.PT.58.19934831), TNF Receptor Superfamily Member 1A (*TNFRSF1A*, Hs.PT.58.19429998) and TNF Receptor Superfamily Member 1B (*TNFRSF1B*, Hs.PT.58.40638488) (all from Integrated DNA Technologies, Coralville IA) according to manufacturer instructions for qRT-PCR. qRT-PCR was performed by the Mount Sinai qPCR CoRE using an ABI PRISM 7900HT sequence detection system with robotic arms (Applied Biosystems, Foster City, CA). Data was analyzed using the 2^−ΔΔCt^ method (77), normalized to housekeeping gene glyceraldehyde 3-phosphate dehydrogenase (GAPDH) and relative to hAF cells cultured basal media. 2^−ΔΔCt^ calculations were conducted using Microsoft Excel (Microsoft, Redmond, NM).

### Bulk RNA sequencing and analyses

Bulk RNAseq was used to evaluate changes in gene expression of hAF cells after TNFα signaling modulation (“TNFR blocking”). RNA was extracted from samples using the RNeasy Micro Kit (QIAGEN INC, USA) according to manufacturer instructions with DNase application and eluted in 15 µL RNase free water. RNA concentration was quantified using a Thermo Scientific NanoDrop™ One UV–Vis spectrophotometer (Thermo Fisher Scientific). RNA integrity (RIN) was measured with an Agilent 2100 BioAnalyzer. 2 μg of total RNA from each sample were used for library preparation. Each library was constructed from RNA from hAF cells from 5 individual donors. First, polyadenylated transcripts were enriched using NEXTFLEX Poly(a) Beads 2.0 (PerkinElmer); then, RNA libraries were prepared using the NEXTFLEX® Rapid Directional RNA-Seq Kit 2.0 (PerkinElmer). Library concentration was measure by Qubit, while library quality and size were calculated with an Agilent 2100 BioAnalyzer. Single-end read sequencing was performed using an Illumina NextSeq 550 instrument and a High-output kit, at a depth of 33 million reads per sample.

Differential expression of bulk transcriptomic analysis involved several steps. FASTQ files of sequenced reads were aligned to human transcriptome (GENCODE annotation model p10 release 27) using the STAR aligner (v2.5.3a), with an average alignment rate of over 80%. Gene-level expression was quantified by the feature Counts (v1.6.3) and normalized by the voom function in R/limma packages. Differential expression was called by R/limma. PCA and clustering were performed in R. Significant DEGs were determined using R/limma at a false discovery rate of 5%, with a fold change greater than or equal to 2. Gene ontology and pathway enrichment was performed using Enrichr to determine significantly enriched or depleted pathways, and normalized enrichment scores reported for top gene ontology and pathway sets.

Network analyses used QIAGEN Ingenuity Pathway Analysis (IPA) (QIAGEN Inc., https://digitalinsights.qiagen.com/IPA) using their developed algorithms (78). Included were DEGs with a FC <-1.5 and >1.5 and a p-value ≤0.01.

Heatmaps for cell cycle were created using bulk RNS-seq results and selecting genes recommended for cell cycle analyses with sequencing data that are relevant across multiple cellular models across different biological contexts (49, 50).

### Senescence and cell cycle analyses

SA-β-Gal staining was performed using the Senescence Detection Kit (Abcam, ab65351) following the manufacturer’s protocol. Briefly, hAF cells were fixed for 15 min with 1× fixation solution, washed with PBS and incubated overnight at 37 °C in 1× staining solution. Brightfield images were captured at 20X on the VWR® Trinocular Inverted Microscope. SA-β-Gal positive cells from 5 biological replicates with n=9 technical replicates per group were analyzed using the ImageJ Cell Counter Plugin.

Cleaved-Casp3 staining involved hAF cells fixed in Z-fix formalin fixative for 10 min at room temperature followed by permeabilization for permeabilization using 0.5% TritonX-100 for 5 minutes. Cells were incubated for 1 hour at room temperature with serum-free protein block (Dako X0909) to block non-specific binding sites followed by Cleaved-Casp3 primary antibody (Abcam, ab49822, 1:1000) at 4°C over night. Next, the secondary antibody (Cy™5 Donkey Anti-Rabbit IgG, 1:400) (Jackson ImmunoResearch, 711-175-152) was added for 1 hour. Finally, sections were cover slipped using ProLong™ Gold antifade reagent with DAPI (Invitrogen P36931). Dual-channel images were captured at 20X on the Zeiss AxioImager fluorescence microscope. For Co-localization analysis, ImageJ Plugin “EzColocalization” was used to quantify Pearson’s correlation coefficient (79). For nuclear fluorescent intensity, the Intensity Density (IntDen) of Casp3 signal within the nuclear reason was determined using ImageJ. A total of 5 biological replicates with n=4 technical replicates were analyzed.

### IHC analysis of human IVDs

Human specimens were fixed in formaldehyde, embedded in resin, and sectioned in the sagittal plane as previously described (80). Mid-sagittal 5°µm sections from each sample were stained with picrosirius red/alcian blue (PR/AB) for morphological reference. Prior to staining, tissue sections were deplasticized and rehydrated. Sections were then treated with proteinase K (S3020, Dako North America, Carpinteria, CA) followed by protein block (X0909, Dako North America). At room temperature, sections were incubated for one hour with either rabbit polyclonal primary antibody against TNF Receptor 1 (dilution 1:400, ab111119, Abcam, Cambridge, MA), recombinant rabbit monoclonal primary antibody against TNF Receptor 2 [EPR1653] (dilution 1:200, ab109322, Abcam, Cambridge, MA) or normal rabbit serum (ab7487, Abcam) as a negative control. Then sections were incubated for 30 minutes at room temperature with anti-rabbit IgG peroxidase secondary antibodies (ImmPRESS reagent, Vector Laboratories, Burlingame, CA). Afterwards the sections were treated with hydrogen peroxide block (ab94710, Abcam) for 10 min followed by diaminobenzidine-based peroxidase substrate for a minute (ImmPACT DAB Vector Laboratories). Finally, the sections were counterstained with 0.1% toluidine blue before mounting. Whole digital slide images were obtained using NanoZoomer S210 from Hamamatsu. Quantification of TNFR1/2 was performed using HALO Image Analysis Platform (v3.6.4134.137, Indica Labs, Albuquerque, NM, USA). Briefly, tissues were annotated to identify the regions to include in the analysis. Once the analysis area is defined, TNFR1/2 is quantified using HALO Multiplex module. The bright filed algorithm uses color deconvolution to separate the chromogenic stains and this allows for the generation of the "markup" image of the of the separated chromogenic stains based on selected settings that will generate the results.

### Statistical analyses

The statistical analyses were performed in R (v4.2.3) using R-based package Seurat (v4.3) for analysis of scRNA-seq data and R-based Limma for analysis of bulk RNA-seq data. For all other data, statistical analyses were performed using GraphPad Prism (v10) software. One-way ANOVA or Two-way ANOVA with Tukey’s post-tests were used to compare between groups. Results were displayed as mean ± standard deviation. Significance was accepted at a level of p < 0.05.

## ACKNOWLEDGEMENTS

We thank Monica Garcia-barros and the Biorepository and Pathology CoRE at the MSSM for the assistance and contribution of the immunohistochemistry quantification using the HALO software. We thank Dr. Nadine Schrode for her assistance and insights on the scRNA-seq analysis. We thank Dr. Nilsson Holguin for useful discussions on interpretations.

## Author contributions

Conceptualization: JCI, JG Study design: JG, EG, JCI Resource acquisition: SC, ACH, WF, CL Performed experiments: JG, EG, DL Analyzed and interpreted the results, and prepared the figures: JG, EG, LR, MW, RS, CL, JCI Funding acquisition: JCI Project administration: JG, JCI Supervision: JCI, JG, RS Writing—original draft: JG, JCI Writing—review, editing and approval: JG, EG, LR, MW, DL, SC, ACH, RS, CL, JCI

**Competing interests:** Authors declare that they have no competing interests.

**Data and materials availability:** All data are available in the main text or the supplementary materials. Data and code for RNA-seq analysis will be made available upon manuscript acceptance.

