## Supplemental Figures and Tables for "TNFR1-mediated senescence and lack of TNFR2-signaling limit human intervertebral disc cell repair in back pain conditions"

### Supplementary Figures

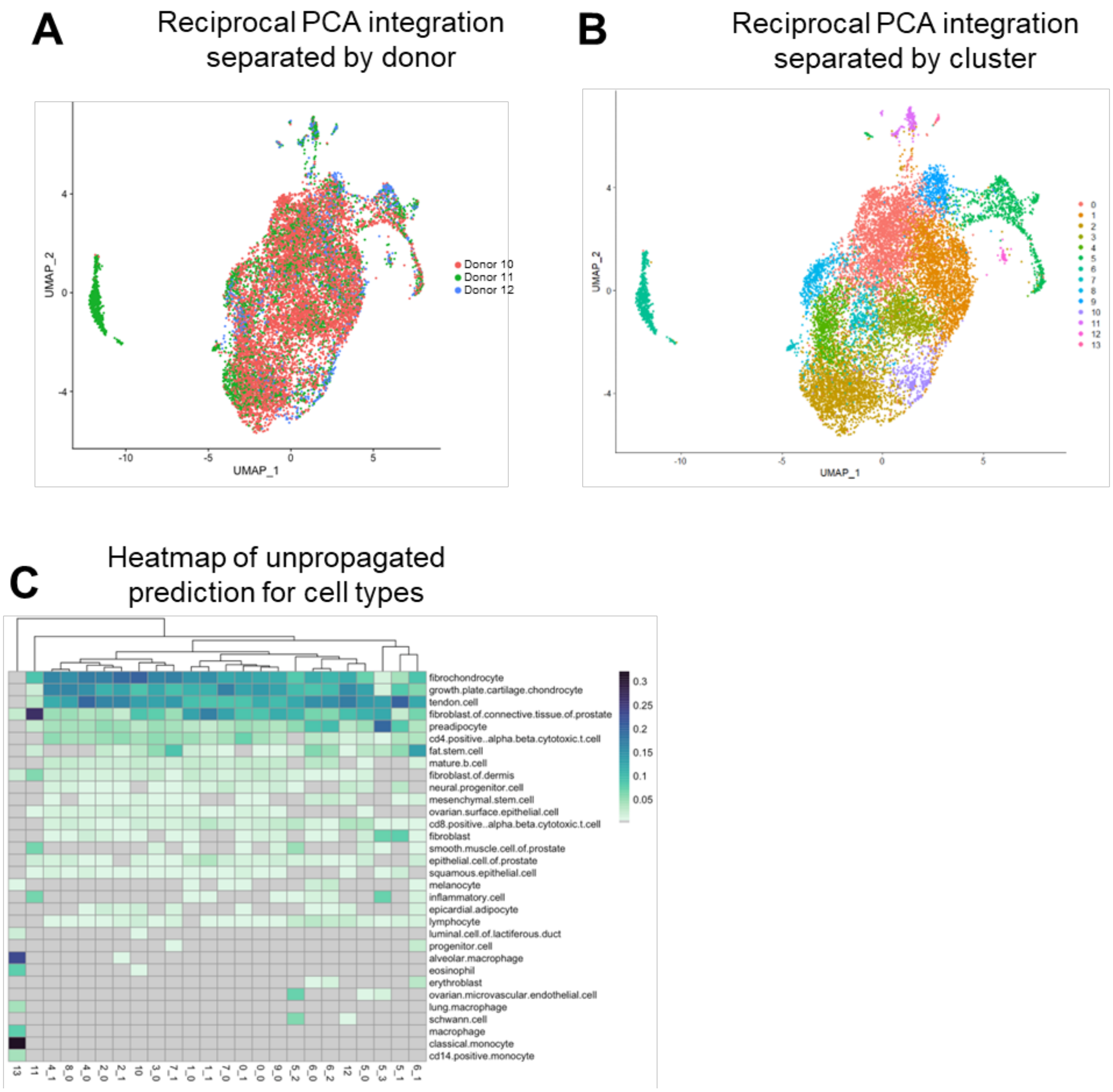


**Fig. S1. scRNA-seq analysis of 13259 cells from 3 biological replicates of human painful IVD tissue specimen in (A)** integrated UMAP, which resulted in **(B)** 14 clusters. **(C)** Unpropagated prediction of cell types for each cluster cell types were annotated using UniCell Deconvolve, a pre-trained, unbiased, deep learning model.

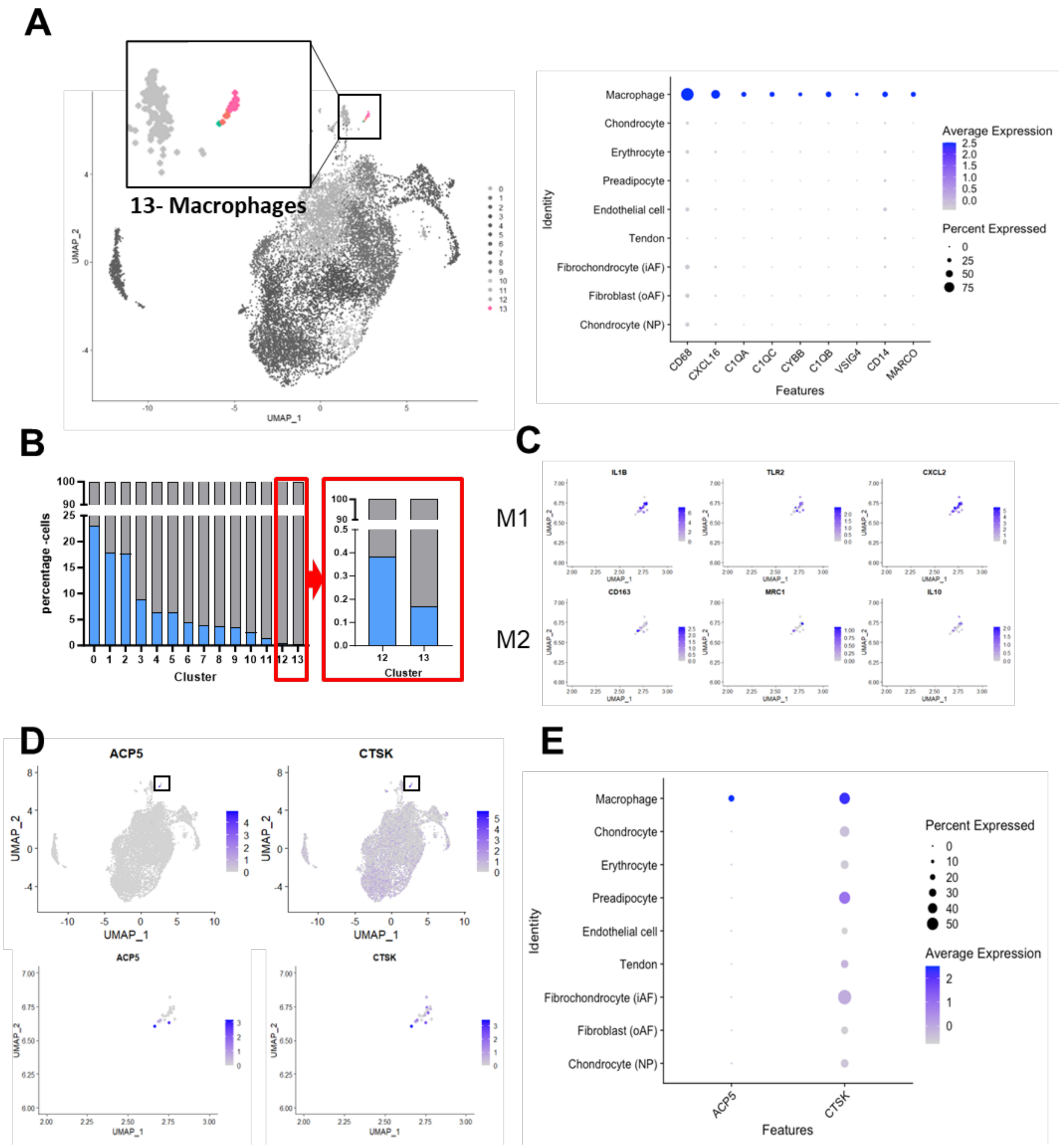


**Fig. S2. Characterization of macrophage cluster 13 was confirmed as (A)** macrophages comparing macrophage markers across all cell population. **(B)** 0.16% of cells withing the herniated IVD belongs to macrophage – cluster 13, of which **(C)** the majority is polarized towards the M1, pro-inflammatory phenotype. **(D and E)** show osteoclast markers present in macrophage population, indication an osteoclast origin of macrophages.

**A**

| Gene | Gene Symbol | Assay ID |
|---|---|---|
| Glyceraldehyde 3-Phosphate Dehydrogenase | GAPDH | Hs.PT.39a.22214836 |
| Collagen Type I Alpha 1 Chain | COL1A1 | Hs.PT.58.15517795 |
| Aggrecan | ACAN | Hs.PT.56a.742783 |
| Matrix Metallopeptidase 13 | MMP13 | Hs.PT.58.40735012 |
| ADAM Metallopeptidase with Thrombospondin Type 1 Motif 4 | ADAMTS4 | Hs.PT.58.19934831 |
| Interleukin 1 Beta | IL1B | Hs.PT.58.1518186 |
| Interleukin 6 | IL-6 | Hs.PT.58.40226675 |
| Tumor Necrosis Factor | TNF | Hs.PT.58.45380900 |

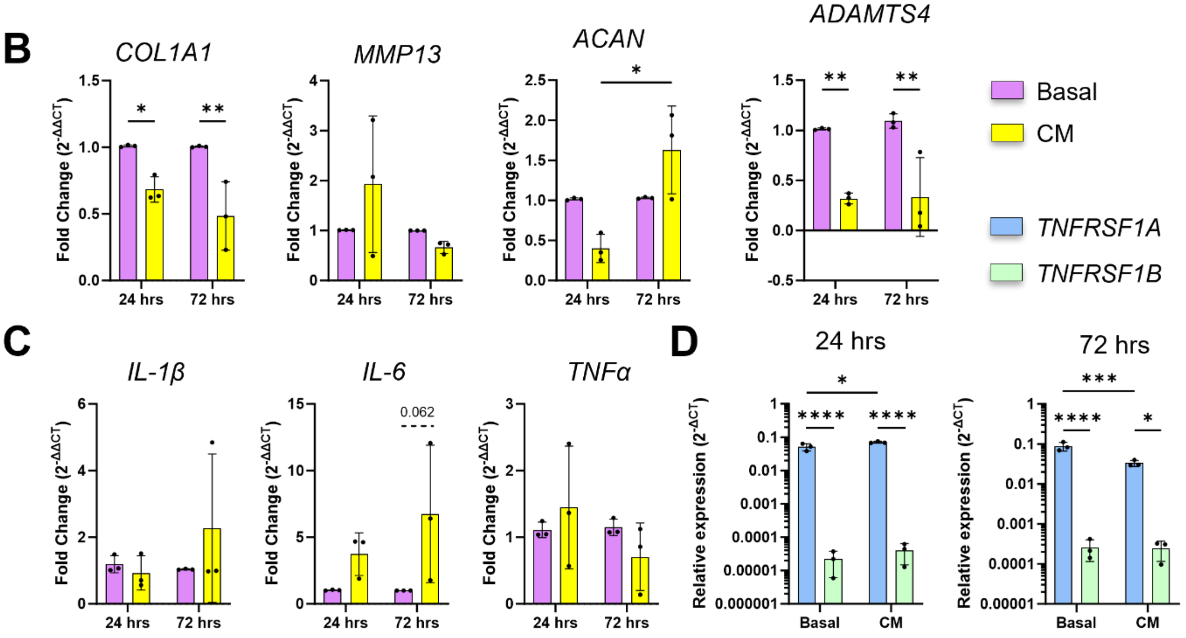


**Fig. S3. hAF cell Gene expression of matrix metabolic and pro-inflammatory genes after 24 and 72 hours in CM (A)** primers being used for qRT-PCR. **(B)** Change in expression of matrix-related genes *COL1A1*, *MMP13*, *ACAN* and *ADAMTS4*. **(C)** Expression of po-inflammatory genes *IL-1β, IL-6* and *TNFα*. (D) Relative gene expression normalized to *GAPDH* of *TNFRSF1A* and *TNFRSP1B* after 24 and 72 hours ($2^{-\Delta CT}$). *, **, *** and **** represent significant differences between groups with p< 0.05, 0.01, 0.001 and 0.0001 respectively.

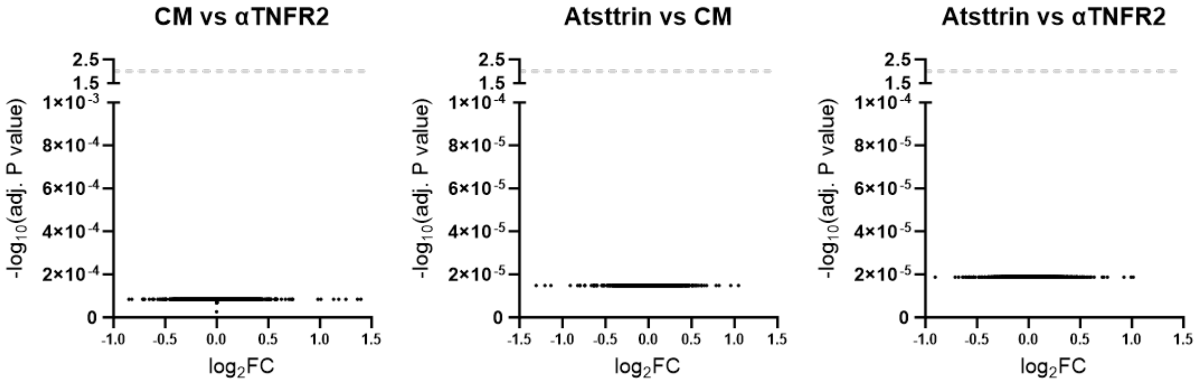


**Fig. S4. Volcano Plots comparing DEGs between CM, αTNFR2 and Atsttrin.** No significant DEGs were found between those groups.

| Donor | Thompson Degeneration Grade | Rutgers Degeneration Grade | TNFR1 pos. cells (%) | | | TNFR2 pos. cells (%) | | |
|---|---|---|---|---|---|---|---|---|
| | | | oAF | iAF | NP | oAF | iAF | NP |
| **17** | 1 | 1.3 | 5.610589 | 6.815001 | 6.869662 | - | - | - |
| **18** | 1 | 3 | 4.470972 | - | - | 7.890606 | 9.582135 | 5.097009 |
| **19** | 2 | 3 | 17.429491 | 11.2069 | - | 13.068454 | 13.02622 | - |
| **20** | 2 | 6 | 4.470972 | - | - | 2.975194 | 62.14198 | - |
| **21** | 2 | 2.3 | 2.943213 | 1.860815 | 0.892857 | - | - | - |
| **22** | 3 | 2.7 | 1.711768 | 1.799527 | 6.930248 | 1.988636 | 2.394184 | 2.113274 |
| **23** | 3 | 9.7 | 16.481943 | 11.64021 | 13.60344 | 1.185609 | 2.459114 | 2.825923 |
| **24** | 3 | 7 | 21.483355 | 11.58234 | 5.008255 | 13.533676 | 6.656761 | 5.890944 |
| **25** | 4 | 8.3 | 9.151707 | 11.03448 | 8.80734 | 4.773786 | 6.882344 | 5.376344 |
| **26** | 4 | 5.3 | 6.482419 | 4.95776 | 13.69095 | 5.982281 | 3.480233 | 2.569078 |
| **27** | 4 | 7.3 | - | - | - | 2.118624 | 2.065782 | 10.22442 |
| **28** | 5 | 5 | 3.572794 | 0.70505 | 0.528159 | 4.049844 | 2.439644 | 1.699926 |
| **29** | 5 | 12 | 14.291757 | 19.45701 | - | 6.112011 | - | - |
| **30** | 5 | 12 | 21.55452 | 31.13592 | 59.0035 | 10.732406 | 22.41131 | - |

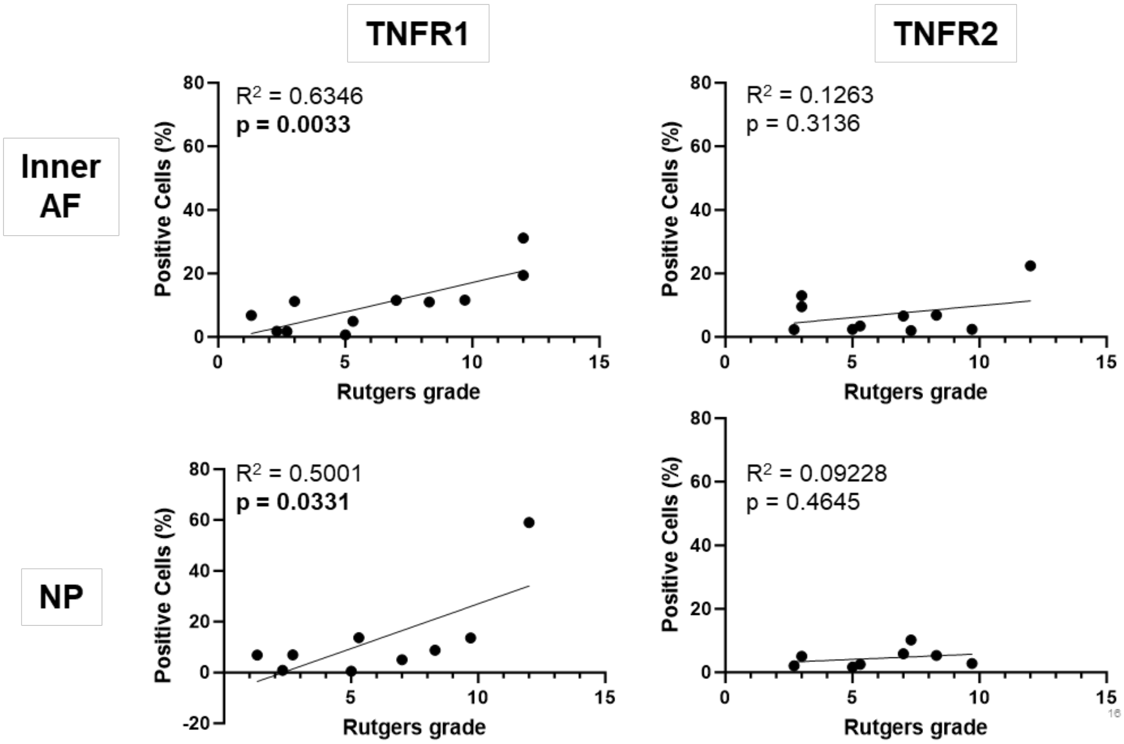




**Fig. S5. Correlation of TNFR1 and TNFR2 to Rutgers degeneration grade in inner AF and NP region of IVD.** There is a significant correlation between TNFR1 and inner AF and NP (p=0.003 and p=0.0331 respectively), which is not observed for TNFR2.

**Table S1.** Demographics of human subject specimens used for in vitro experiments.

| Donor | Age | Sex | Diagnosis | Pfirrrman Degeneration Grade | Region | Media Conditioning | scRNAseq | Timing | Dosing | TNFR blocking |
|---|---|---|---|---|---|---|---|---|---|---|
| **1** | 34 | F | Disc herniation | 2 | Cervical | AF +NP | | AF | AF | AF |
| **2** | 45 | M | Disc herniation | 4 | Lumbar | AF +NP | | AF | AF | AF |
| **3** | 47 | M | Disc herniation | 2 | Cervical | AF +NP | | | | |
| **4** | 49 | M | Disc herniation | 3 | Cervical | AF +NP | | | | |
| **5** | 50 | M | Disc herniation | 3 | Cervical | AF +NP | | | | |
| **6** | 53 | M | Disc herniation | 2 | Cervical | AF +NP | | | | |
| **7** | 55 | F | Disc herniation | 4 | Cervical | AF +NP | | | | |
| **8** | 56 | F | Disc herniation | 3 | Cervical | AF +NP | | | | |
| **9** | 65 | M | Disc herniation | 5 | Cervical | AF +NP | | | | |
| **10** | 52 | M | Disc Degeneration and herniation | 4 | 2x Cervical | | AF +NP | | | |
| **11** | 37 | F | Disc herniation | 2 | Cervical | | AF +NP | | | |
| **12** | 67 | F | Disc Degeneration and herniation | 4 | 2x Cervical | | AF +NP | | | |
| **13** | 25 | M | Disc herniation | 3 | Cervical | | | | AF | AF |
| **14** | 47 | F | Disc herniation | 4 | Cervical | | | | AF | AF |
| **15** | 53 | M | Disc herniation | 3 | Cervical | | | AF | AF | |
| **16** | 71 | M | Disc herniation | 4 | Cervical | | | | AF | AF |

**Table S2.** 44 inflammation-related cytokines found in CM and Basal media (in pg/mL +/- SD)

| Factor | Concentration (pg/mL ± SD) Basal | CM | Basal vs CM P-value | Sig |
|---|---|---|---|---|
| sCD40L | 0.19 ± 0.61 | 3.81 ± 3.14 | 0.0006 | *** |
| EGF | 3.031 ± 0.33 | 4.14 ± 1.35 | 0.0160 | * |
| Eotaxin | 0.58 ± 0.52 | 10.49 ± 7.75 | 0.0001 | *** |
| FGF-2 | 6225.60 ± 749.42 | 794.76 ± 561.81 | <0.0001 | **** |
| FLT-3L | 0 ± 0 | 1.06 ± 0.61 | <0.0001 | **** |
| Fractalkine | 2.42 ± 3.93 | 10.93 ± 5.60 | 0.0004 | *** |
| G-CSF | 0 ± 0 | 5.22 ± 7.94 | 0.0491 | * |
| GM-CSF | 0.15 ± 0.46 | 1.12 ± 1.91 | 0.1676 | ns |
| GROα | 0.58 ± 0.20 | 74.01 ± 130.01 | 0.1021 | ns |
| IFN-α2 | 1.35 ± 1.54 | 4.25 ± 1.28 | <0.0001 | **** |
| IFNγ | 0 ± 0 | 0.01 ± 0.21 | 0.1937 | ns |
| IL-1α | 0.35 ± 1.09 | 0.45 ± 1.17 | 0.3646 | ns |
| IL-1β | 1.65 ± 0.60 | 1.96 ± 1.45 | 0.7726 | ns |
| IL-1RA | 0.048 ± 0.14 | 0.34 ± 0.29 | 0.0116 | * |
| IL-2 | 0 ± 0 | 0.06 ± 0.08 | 0.0464 | * |
| IL-4 | 0 ± 0 | 1.05 ± 0.88 | 0.0003 | *** |
| IL-6 | 0 ± 0 | 319.97 ± 334.52 | 0.0032 | ** |
| IL-7 | 0 ± 0 | 0.06 ± 0.17 | 0.4245 | ns |
| IL-8 (CXCL8) | 0.10 ± 0.09 | 1510.30 ± 1801.72 | 0.0104 | * |
| IL-9 | 0.42 ± 0.18 | 0.92 ± 0.50 | 0.0050 | ** |
| IL-10 | 0 ± 0 | 1.43 ± 1.52 | 0.0040 | ** |
| IL-12p40 | 1.89 ± 1.59 | 3.98 ± 3.08 | 0.0786 | ns |
| IL-12p70 | 0.07 ± 0.21 | 3.52 ± 3.75 | 0.0048 | ** |
| IL-13 | 0.74 ± 0.93 | 33.32 ± 21.44 | <0.0001 | **** |
| IL-15 | 0 ± 0 | 0.54 ± 0.88 | 0.0730 | ns |
| IL-17A | 0 ± 0 | 0.17 ± 0.30 | 0.1267 | ns |
| IL-18 | 0.07 ± 0.08 | 0.23 ± 0.16 | 0.0093 | ** |
| IL-22 | 0 ± 0 | 20.55 ± 14.68 | <0.0001 | **** |
| IL-27 | 6.27 ± 5.69 | 12.66 ± 5.04 | 0.0397 | * |
| IP-10 | 0.06 ± 0.17 | 704.53 ± 937.84 | 0.0221 | * |
| MCP-1 (CCL2) | 0.05 ± 0.17 | 504.40 ±240.55 | <0.0001 | **** |
| MCP-3 (CCL7) | 0.34 ± 0.72 | 8.68 ± 4.92 | <0.0001 | **** |
| M-CSF | 0 ± 0 | 172.1 ± 68.62 | <0.0001 | **** |
| MDC | 0 ± 0 | 118.12 ± 70.66 | <0.0001 | **** |
| MIG/CXCL9 | 0 ± 0 | 997.98 ± 761.82 | <0.0001 | **** |
| MIP-1α (CCL3) | 0.08 ± 0.24 | 7.16 ± 17.33 | 0.2935 | ns |
| MIP-1β (CCL4) | 1.02 ± 0.63 | 5.47 ± 7.15 | 0.0703 | ns |
| PDGF-AA | 0 ± 0 | 107.76 ± 69.30 | <0.0001 | **** |
| PDGF-AB/BB | 0.97 ± 3.06 | 2.24 ± 4.47 | 0.6291 | ns |
| RANTES (CCL5) | 0.06 ± 0.12 | 91.22 ± 63.69 | <0.0001 | **** |
| TGFα | 0.13 ± 0.19 | 4.08 ± 3.05 | 0.0001 | *** |
| TNFα | 0 ± 0 | 3.91 ± 2.43 | <0.0001 | **** |
| TNFβ | 0.08 ± 0.17 | 14.21 ± 10.86 | <0.0001 | **** |
| VEGF-A | 0 ± 0 | 550.90 ± 362.30 | <0.0001 | **** |

**Table S3.** Demographics of autopsy samples for IHC of TNFR1 & TNFR2

| Donor | Age | Sex | Cause of Death | Thompson Degeneration Grade | Rutgers Degeneration Grade | Level |
|---|---|---|---|---|---|---|
| **17** | 28 | M | liver and renal failure from testicular cancer | 1 | 1.3 | Lumbar |
| **18** | 30 | F | sepsis | 1 | 3 | Lumbar |
| **19** | 22 | F | Liver Failure | 2 | 3 | Lumbar |
| **20** | 54 | M | alcohol liver disease | 2 | 6 | Lumbar |
| **21** | 58 | F | While obese , respirator 3 days, brain dead | 2 | 2.3 | Lumbar |
| **22** | 32 | F | liver failure | 3 | 2.7 | Lumbar |
| **23** | 40 | M | Lung infection | 3 | 9.7 | Lumbar |
| **24** | 52 | M | Melanoma Metastasis | 3 | 7 | Lumbar |
| **25** | 47 | F | Cancer | 4 | 8.3 | Lumbar |
| **26** | 48 | M | Huntingtons disease Respiration | 4 | 5.3 | Lumbar |
| **27** | 66 | F | n/a | 4 | 7.3 | Lumbar |
| **28** | 62 | F | Heart Disease | 5 | 5 | Lumbar |
| **29** | 85 | M | Respiratory Failure/ Heart Failure/Hyptertension | 5 | 12 | Lumbar |
| **30** | 85 | M | Parkinsons + Arthrosclerotic disease | 5 | 12 | Lumbar |
